# A nucleic acid labeling chemistry reveals surface DNA on exosomes

**DOI:** 10.1101/2025.11.18.689134

**Authors:** Filip Boskovic, Priyanka Dutta Gupta, Jian Zhang, Yamuna Krishnan, Jack W. Szostak

**Affiliations:** Howard Hughes Medical Institute, University of Chicago, Chicago, IL, 60637, USA; Department of Chemistry, University of Chicago, Chicago, IL, 60637, USA; Neuroscience Institute, The University of Chicago, Chicago, IL, 60637, USA; Institute for Biophysical Dynamics, The University of Chicago, Chicago, IL, 60637, USA

**Keywords:** nucleic acid chemical labeling, nucleic acid chemistry, surface DNA, exosomes

## Abstract

Chemical labeling of nucleic acids is essential to pinpoint the structure, localization, and function of RNA and DNA. Yet, reversible sequence-independent chemistries that can label native RNA and DNA remain poorly developed. Here we describe Reversible Uridine Nitrilium-mediated Addition (RUNA), a reversible covalent chemistry that selectively modifies uridine and thymidine residues *via* N3 deprotonation and reaction with a nitrilium ion intermediate generated from an aldehyde and an isonitrile. The reaction forms a stable N3 adduct that can be quantitatively reversed by hydrolysis. By using reagents that are either membrane permeable or impermeable, we demonstrate the localization and function of DNA on exosomes. Although exosomes harbor nucleic acids, whether the latter are encapsulated in the exosome lumen or are surface-adhered is unknown. RUNA revealed that exosomes display DNA on their outer surface. The abundance of such surface DNA increases upon DNA-damage accumulation in cancer cells that are treated with a PARP inhibitor. This surface DNA drives exosome uptake by M2-polarized macrophages through scavenger receptors and triggers a shift toward an M1-like pro-inflammatory state. The selective labeling of surface DNA revealed an unexpected mechanism by which exosomes engage innate immune cells. RUNA is a versatile tool to analyze the nucleic acid content and functionality of extracellular vesicles in health and disease.

**Significance Statement:** Pinpointing the localization of RNA and DNA in cells and organelles is central to deriving insights into their biological functions in health and disease. We describe a new method, RUNA, for labeling nucleic acids that is sequence-independent and reversible. By varying RUNA reagents, we can distinguish between nucleic acids that are located either inside or outside of membrane compartments. Using RUNA, we showed that DNA is associated with the outer surface of exosomes that are secreted by cancer cells. Further, the amount of surface DNA increases when the cancer cells are treated with an anti-cancer drug. This surface DNA promotes the uptake of exosomes by innate immune cells known as macrophages and modulates their inflammatory response.

## Introduction

The introduction of a label on RNA or DNA, independent of sequence, requires the functionalization of either the nucleobase or the sugar-phosphate backbone (1–4). Such labels can include fluorophores for imaging, affinity handles for enrichment and purification, or reactive groups for bioorthogonal conjugation (5–9). Such tags enable the study of nucleic acid composition, structure, localization, and function in biology (10–14). While this is best achieved through enzymatic strategies that are minimally disruptive, enzymatic methods are generally sequence specific (15–17). For the unbiased analysis of nucleic acids, sequence-independent labeling chemistries are highly desirable. For example, RNA labeling by 2′-hydroxyl acylation or sulfonylation enables the mapping of RNA structures and RNA-RNA interactions (1, 18). Nucleobase-specific chemistries using glyoxal and kethoxal that selectively label unpaired guanosines (19, 20) led to the development of sequencing methods such as KAS-seq or KARR-seq that are used to map transcriptionally-active DNA loci (21) or RNA-RNA interactions, respectively (2). Carbodiimide reagents such as 1-ethyl-3-(3-dimethylamino-propyl)carbodiimide (EDC) and N-cyclohexyl-N′-(2-morpholinoethyl)carbodiimide (CMC), which modify uridine and guanosine, form the basis of pseudouridine sequencing (4, 22).

New biological roles are emerging for membrane-associated nucleic acids in cellular communication and immune regulation (23–26). The importance of surface RNA highlights the need for mild, sequence-independent labeling strategies that can help to define RNA location, synthesis and functionality (26–28). GlycoRNAs represent such a class, given their emerging roles in receptor engagement (29) and immune signaling (23–25). Circulating tumor DNA has gained importance in cancer diagnosis (28, 30). Both tumor cells and normal cells release small extracellular vesicles, including exosomes, that are associated with nucleic acids (23, 27, 30). However, it is unclear whether the DNA is inside the exosome or is surface associated (27). Although this distinction is important for understanding how exosomes mediate immune responses, it has remained unresolved because there is no way to chemically discriminate DNA molecules on the inside of exosomes from those on the outside.

To probe nucleic acids in such cell-derived compartments sequence-independently, we developed a reversible labeling chemistry that selectively modifies uridines and thymidines in RNA and DNA respectively. It couples an aldehyde and an isonitrile to generate a nitrilium ion *in situ*, which reacts specifically at the N3 position of uracil and thymine to form a stable covalent adduct. This reaction, termed Reversible Uridine Nitrilium-mediated Addition (RUNA), is highly modular. By changing the aldehyde, one can install diverse functional groups on the DNA or RNA. If the tag is membrane-impermeant, one can probe nucleic acid accessibility across biological membranes.

We applied RUNA to resolve the long-standing conundrum of whether exosomal DNA is entrapped within the lumen or adhered to the outer surface (27, 30–32). By applying RUNA to exosomes derived from prostate cancer cells, we found that DNA is present on the surface of exosomes. When the prostate cancer cells are treated with a poly(ADP-ribose) polymerase (PARP) inhibitor such as rucaparib, RUNA indicates that the abundance of surface-attached DNA increases, revealing that DNA damage can influence the amount of extracellular DNA. Importantly, such exosomes are more efficiently internalized by M2-polarized macrophages, which then adopt a more M1-like macrophage state and a proinflammatory cytokine response. These results reveal exosomal surface DNA as both a determinant of exosome uptake and a modulator of macrophage phenotype. Our studies position RUNA as a versatile chemical platform for the reversible labeling of nucleic acids with spatial resolution across membranes. Our approach opens new avenues to explore the composition and function of membrane-associated RNA and DNA in physiology and disease.

## Results

### Mechanism of native RNA and DNA labeling with RUNA chemistry

Here, we describe RUNA, a uridine- and thymidine-selective chemistry that labels nucleic acids reversibly and sequence-independently. RUNA proceeds *via* the *in situ* generation of a nitrilium ion intermediate. The reaction begins with the condensation of an aldehyde and an isonitrile to form a nitrilium intermediate, which is then attacked by the deprotonated N3 of uridine or thymidine, yielding a covalent adduct (33, 34). Because this modification is thermally labile it can be hydrolyzed by mild heating to restore the native nucleobase (**Scheme 1**). Thus, RUNA can be used to reversibly tag both RNA and DNA with a variety of functional groups such as alkynes, alkenes, and strained cycloalkenes (**Figure 1a**).

**Scheme 1.**
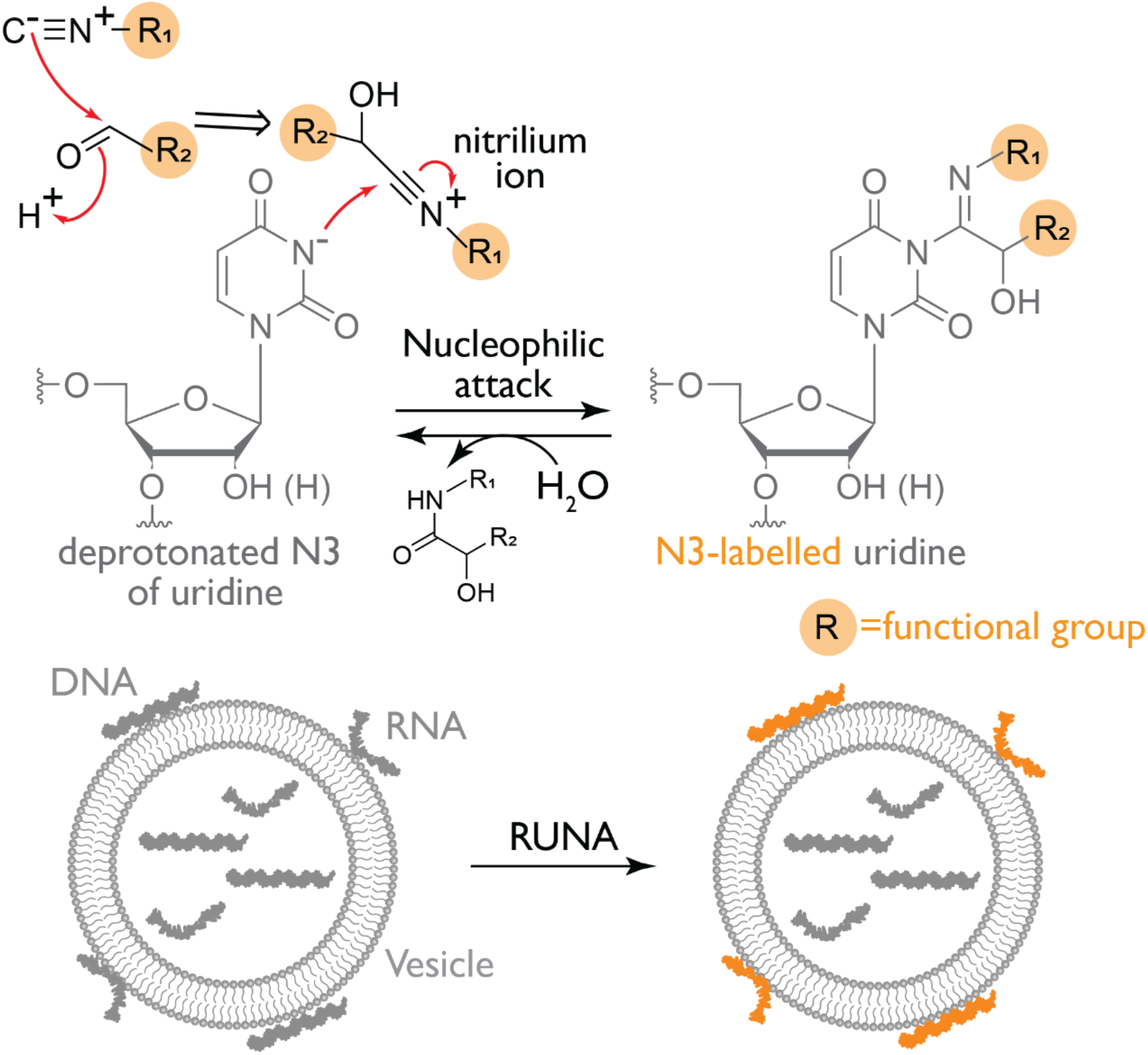
The proposed mechanism for the uridine-preferred labeling of N3 *via* an *in situ* formed nitrilium ion. The deprotonated N3 of uridine (or thymidine) acts as a nucleophile, attacking the nitrilium ion generated from the reaction of an aldehyde with an isonitrile (where R represents a functional group of interest). The resulting modification is hydrolyzed on demand by heating, thereby restoring the native RNA/DNA. The choice of functional groups enables membrane-specific labeling, allowing selective modification of extravesicular or surface-exposed nucleic acids.

**Figure 1.**
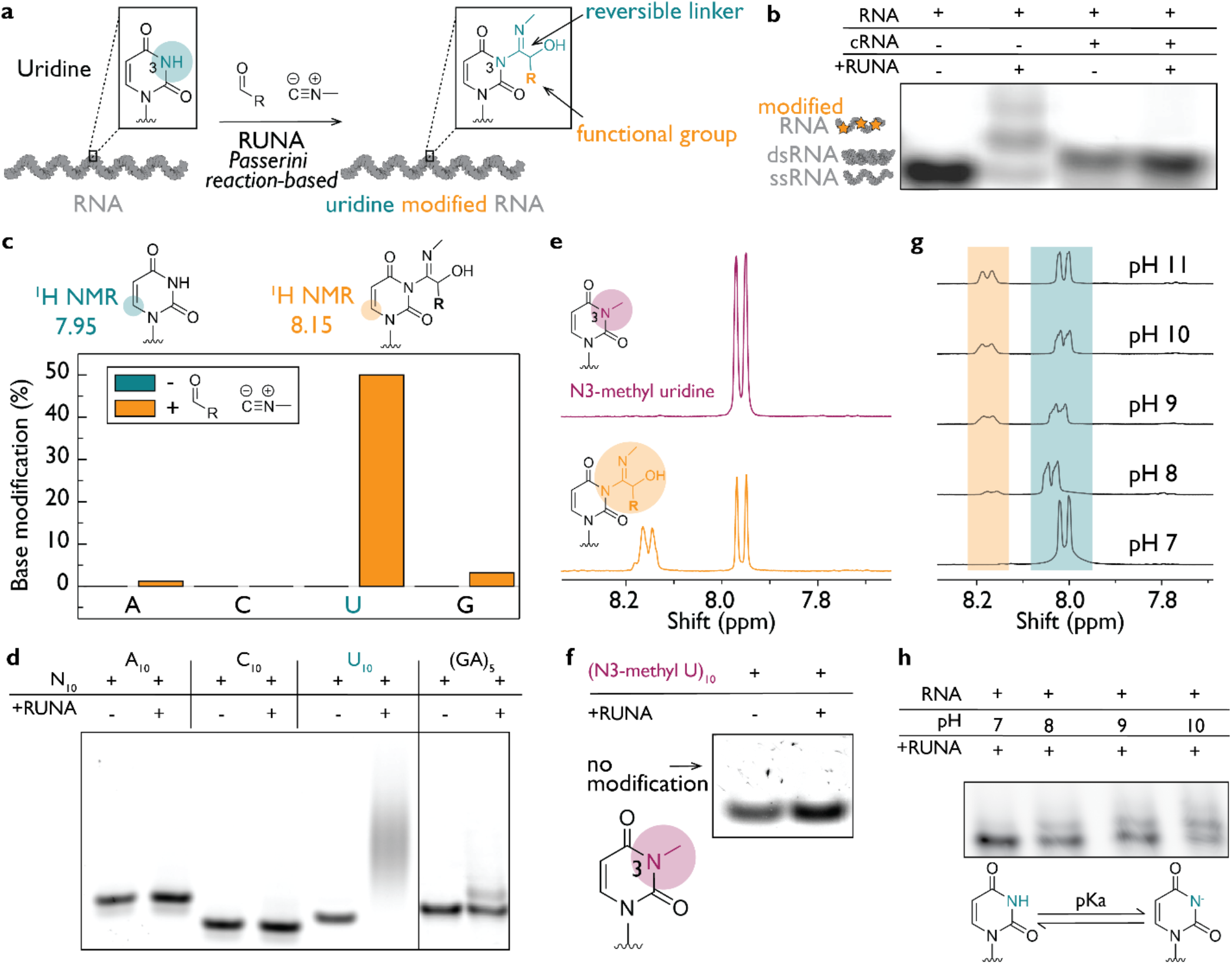
Nucleotide-specific RNA labeling involves N3 of uridine. (**a**) Uridine in RNA is modified to form an N3 adduct containing a removable linker and a functional group of interest. (**b**) Gel analysis shows that single-stranded RNA (12-nt Cy3-labeled) reacts with RUNA chemistry, while double-stranded RNA made by mixing RNA and complementary RNA (cRNA) is not modified. (**c**) The extent of N3 adduct formation as quantified by the proton shift observed in the ^1^H NMR spectrum. Uridine (U) is modified at a higher rate than other nucleotides. (**d**) 10-mer oligonucleotides with a 3′ Cy3 label indicate preferential modification of U_10_ over other oligonucleotides. (**e**) Modification occurs exclusively on U by comparing modification of N3-methylU (purple) with U (orange) under identical conditions. (**f**) No modification occurs if the N3 position is inaccessible as see with an N3-methylU oligonucleotide labeled with 3′ Cy3. (**g**) The pH dependence of labeling indicates that deprotonation of N3 is responsible for modification. (**h**) Gel analysis as a function of reaction pH. For all panels, 4-pentenal and methyl isonitrile were used as the aldehyde and isonitrile respectively.

We first sought to determine whether base pairing of uridine affects its reactivity to RUNA, employing methyl isonitrile (MeNC) and 4-pentenal. We compared the labeling efficiency of single-stranded versus double-stranded RNA (**Figure 1b**). Using denaturing polyacrylamide gel electrophoresis (PAGE), we observed that RUNA labels single-stranded RNA (lane 2) to a greater extent than duplex RNA (lane 4). Interestingly, RUNA labels both single-stranded and duplex DNA with high efficiency (**Figure S1**). The difference in reactivity between dsDNA and dsRNA is most likely due to the higher stability of the dsRNA duplex.

To determine nucleobase selectivity, we monitored RUNA labeling on the four canonical ribonucleotides using ^1^H nuclear magnetic resonance (NMR) spectroscopy. Only uridine monophosphate (UMP) showed a new diagnostic proton signal upon modification, consistent with the formation of an N3 adduct (**Figure 1c**; **Figures S2–S9**). When we rigorously tested this preference using short oligonucleotides, we found high selectivity for rU_10_, over rA_10_, rC_10_, and r(GA)_5_ (**Figure 1d**; **Table S1**). Further, U_10_ exhibits at least one modification per molecule unlike other oligonucleotides, where labeling was negligible under the same conditions. We observed similar reactivity for deoxyribonucleotides, showing that RUNA selectively targets U and T in RNA and DNA respectively (**Figure S10**).

We hypothesized that RUNA labels the N3 of uridine by deprotonation, followed by nucleophilic attack on the *in situ* generated nitrilium ion to yield the N3 adduct. To test this mechanism, we evaluated both free N3-methyl uridine (N3-Me U) and an N3-Me U oligomer with RUNA, and in both cases found that N3 methylation blocks uridine modification (**Figure 1e-f, Figure S11**). To test the need for N3 deprotonation, we examined the pH dependence of RUNA labeling (**Figure 1g, h**). Higher pH led to higher yields of the N3 adduct, consistent with its pKa of 9.2 (**Figure S12**). We tested a range of aldehydes at pH values from 7 to 10 and found that labeling was generalizable across aldehydes (**Figure S13-S16**). Although many are compatible with RUNA, long-chain aliphatic and highly hydrophobic aldehydes such as C8, C10, and pyrene derivatives do not support efficient modification, primarily due to solubility problems (**Figure S17**).

We next evaluated potential side reactions and tested the scope of isonitriles and aldehydes compatible with RUNA. One possible side reaction involves formation of a 5′-imidoyl phosphate intermediate, which could undergo a Passerini-type rearrangement to yield a stable phosphodiester byproduct (33). However, this byproduct was undetectable (**Figure S18**). The isonitrile moiety must be attached to a primary carbon for efficient nitrilium ion formation, since secondary or tertiary substitutions led to poor yields (**Figure S19**). As an alternative to the volatile methyl isonitrile, we found that a safer alternative, 2-morpholinoethylisonitrile, supported efficient labeling (**Figure S20**). Indeed, this chemistry modifies RNA and DNA with various reactive groups such as alkenes, alkynes, and strained cycloalkenes (**Figures S13-S16**).

Lastly, we found that RUNA can label RNA with norbornene aldehyde, which is amenable to click chemistry, with an observed rate of 0.33 ± 0.01 min^−1^ (**Figure S21**). The resulting norbornene-modified RNA reacted with biotin-PEG_4_-tetrazine through an inverse electron-demand Diels–Alder reaction, thereby tagging the RNA at a rate of 0.44 ± 0.02 min^−1^ (**Figure S22**). In sum, RUNA labeling was remarkably efficient in near-native environments of physiological ion levels, pH and temperature, molecular crowding and low mM reagent concentrations (**Figure S23**).

### Reversible functionalization of native RNA

We next investigated whether the N3 adduct on uridines could be removed to restore native RNA structure (**Figure 2**). We first assessed product stability and found that the introduced modifications remained as such for >7 days at room temperature. To test if the adduct could be removed upon heating, we heated modified UMP and monitored the reaction by ^1^H NMR spectroscopy at pH 8.0 (**Figure 1b**). Upon heating to 95 °C for 15 minutes, the U-N3 adduct was completely converted to native uridine (**Figure 2d**; **Figure S24**).

**Figure 2.**
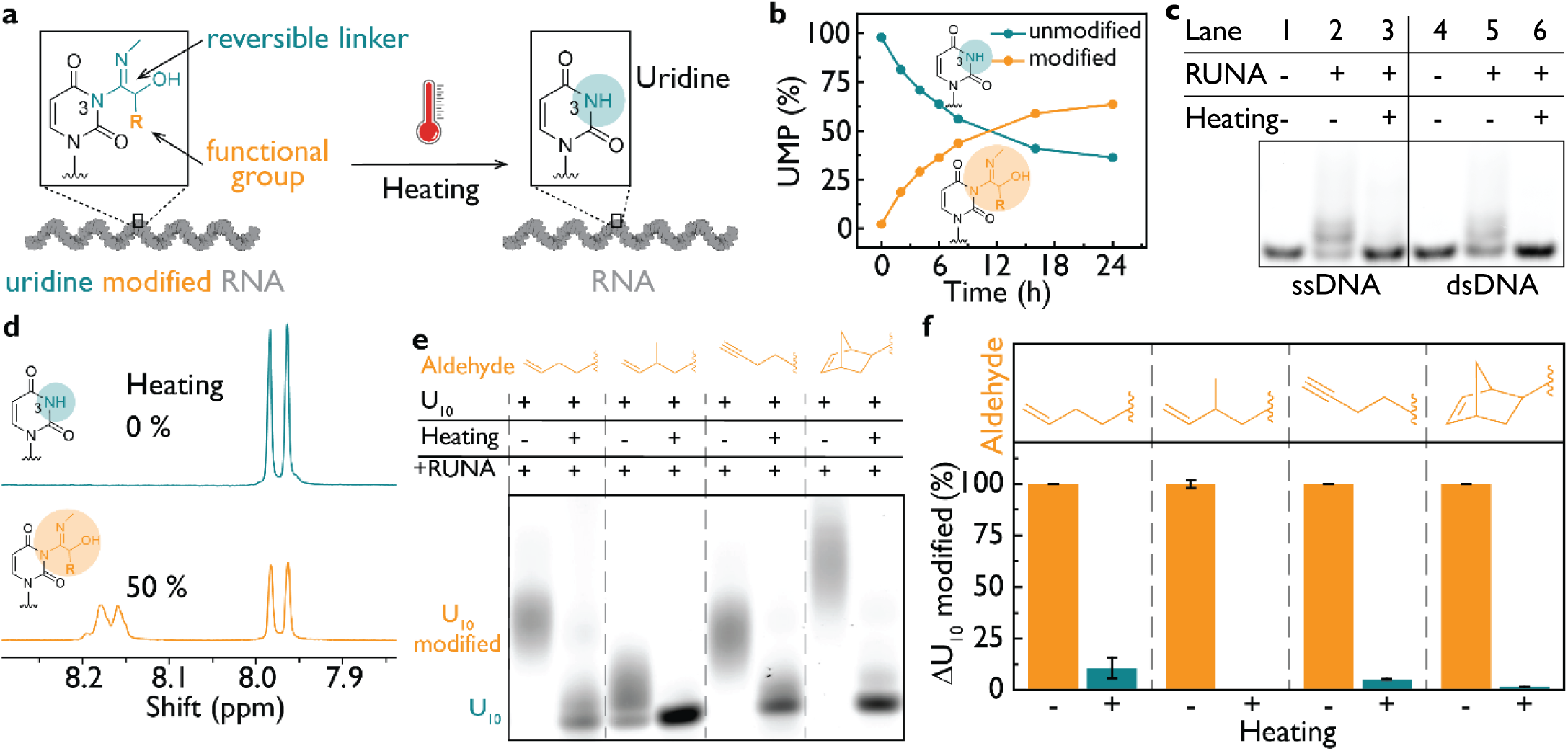
Reversible functionalization of native RNA. (**a**) RNA containing modified uridine is restored to native RNA state by hydrolysis upon heating. (**b**) Time course of UMP modification in the presence of 100 mM MeNC and norbornene aldehyde, pH 8.0, given by ^1^H NMR (**c**) Reversibility of RUNA labeling of ssDNA and dsDNA, analyzed by 20% (v/v) denaturing urea PAGE. Lanes: (1-3) ssDNA; (4-6) dsDNA without or with RUNA treatment (100 mM, 12 h) or after heating at 95 °C for 10 min. (**d**) ^1^H NMR showing that the N3 modification of uridine is removed by heating at 95 °C for 15 min at pH 8.0. (**e**) Reversible modification of U_10_-Cy3 RNA with various functional groups, including tags such as alkene, alkyne, and strained norbornene. (**f**) Reversibility of U_10_ modifications after heating, shown by gel mobility shift analysis.

We next evaluated whether this chemistry also applied to oligonucleotides. We found that RUNA efficiently labeled both single-stranded DNA (ssDNA) and duplex DNA (dsDNA). The labeling was reversed upon heating, as seen by denaturing PAGE (**Table S1, Figure 1c**). We labeled rU_10_ by RUNA with 4-pentenal, 2-methyl-4-pentenal, 4-pentynal, or norbornene aldehyde (5-norbornene-2-carboxaldehyde), and upon heating, we observed efficient adduct removal in all cases (**Figure 2e, f**; **Figure S25**). We demonstrate that RUNA reversibly labels full-length 3.6 kb MS2 RNA; the starting material can be recovered with no detectable loss of integrity (**Figure S26**) under divalent ion–free conditions (35). Because the N3 adduct formed by RUNA is thermally labile, any functional group introduced by this method can be efficiently and cleanly reversed to yield the native RNA.

### Tuning aldehyde membrane permeability selectively labels total versus extravesicular RNA

We then tested whether we could distinguish between intra- and extravesicular RNA by tuning the membrane permeability of the aldehyde reagent. We developed a model system in which phospholipid vesicles composed of 1-palmitoyl-2-oleoyl-sn-glycero-3-phosphocholine (POPC) encapsulated an ATTO647-labeled RNA (blue). A second, extravesicular RNA of identical sequence labeled with ATTO488 (green) was added to the bulk solution (**Figure 3a, Figure S27**).

**Figure 3.**
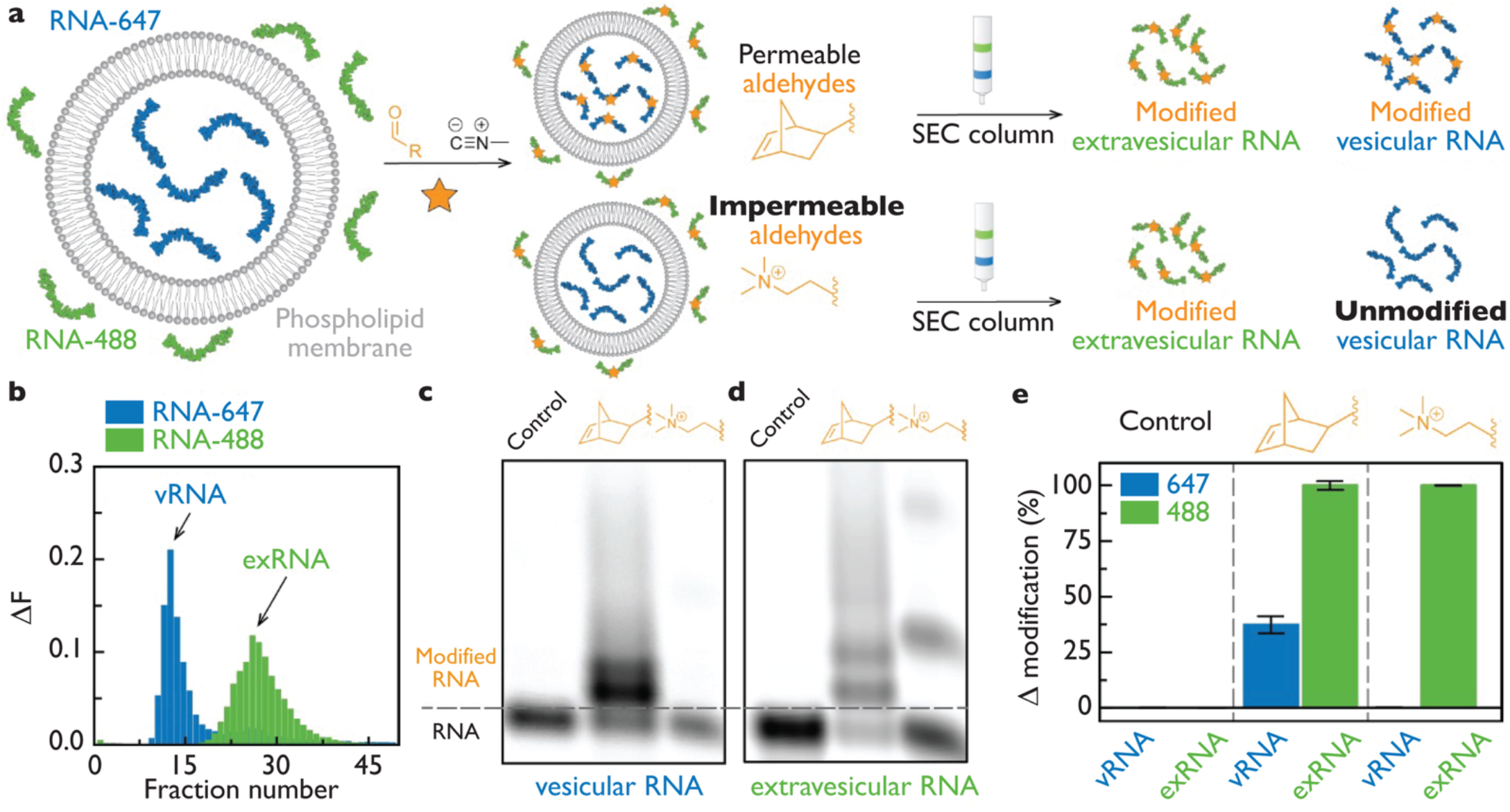
Selective labeling of extravesicular RNA (exRNA) versus vesicular RNA (vRNA) leveraging aldehyde permeability. **(a)** Schematic of selective labeling of exRNA. blue; vRNA; green; exRNA; orange star represents modification. **(b)** Representative fluorescence trace of SEC peaks corresponding to vRNA and exRNA **(c, d)** Denaturing gels of the vRNA and exRNA fractions. Samples with isonitrile but without aldehyde shows no detectable labeling. **(e)** Quantification of RNA modification levels: no labeling without aldehyde, norbornene aldehyde labels both vRNA and exRNA , and betaine aldehyde labels only exRNA.

Membrane-permeable hydrophobic aldehydes can access both RNA pools, whereas charged aldehydes can access only extravesicular RNA. To test whether we could chemically distinguish between both RNA populations, we used a membrane-permeable norbornene aldehyde and a membrane-impermeant betaine aldehyde. Following RUNA labeling, vesicular RNA (vRNA) and extravesicular RNA (exRNA) were separated by size-exclusion chromatography (**Figure S28**). Vesicles containing vRNA eluted early, while exRNA eluted later (**Figure 3b**). Peak fractions corresponding to vRNA and exRNA were collected, and RNA from both fractions was analyzed by denaturing PAGE. Labeling was undetectable without aldehyde (**Figure 3c, d**). The membrane permeable norbornene aldehyde labeled both vRNA and exRNA. In contrast, betaine aldehyde selectively labeled exRNA, consistent with its exclusion from the vesicle interior (**Figure 3e**). Thus, aldehyde permeability can be leveraged to selectively label extravesicular nucleic acids using RUNA.

### RUNA reveals that exosomes carry DNA on their surface

We then sought to test whether RUNA could differentiate between extravesicular and luminal DNA in cell-derived vesicles. Exosomes (Exo) are nanoscale vesicles that cells release into the extracellular environment. They have attracted considerable attention for their burgeoning roles in intercellular communication and in drug delivery. The nucleic acid components of exosomes play key roles in signaling and disease (36). Although transcriptomic and proteomic analyses have catalogued exosomal contents, the precise localization of the nucleic acid components within or on the exosome remains unresolved. To resolve this conundrum, we generated exosomes from a cancer cell line before and after treatment with an anti-cancer therapeutic. We used c-Myc driven murine prostate cancer cells (MyC-CaP) and treated them with 0.5 µM rucaparib (Ruc), a clinically used PARP inhibitor (37) (**Figure 4a**). Because rucaparib treatment impairs DNA repair, induces genomic stress and is known to promote the accumulation of cytosolic DNA (38), we sought to test whether it also affected the DNA content of secreted exosomes. Exosomes released by MyC-CaP cells under either condition were isolated as previously described (39), and characterized in our hands by dynamic light scattering (**Figure S29-S30**) as well as key exosome markers (**Figure S31)**. We found that rucaparib treatment did not alter either the abundance or the size distribution of exosomes produced by MyC-CaP cells (**Figure S29-S30**), suggesting that rucaparib leaves exosome biogenesis largely unaltered. We first probed the DNA content of exosomes by ethidium bromide (EtBr) labeling. When EtBr intercalates into dsDNA, its fluorescence anisotropy increases. Exosomes obtained from cells with or without rucaparib treatment showed an increase in anisotropy, which did not change upon RNase A or proteinase K treatment, reaffirming that the observed signal was DNA-specific (**Figure S32**). When exosomes were treated with DNase I, which should selectively degrade exposed DNA, the anisotropy reduced dramatically (**Figure 4b)**, suggesting that at least some of the DNA associated with exosomes is accessible to DNase 1 and is exposed to the bulk solution, which we denote as surface DNA. Alternatively, the surface DNA may comprise only a small, but structurally critical fraction, and its removal could expose the major lumenal fraction by disrupting exosome integrity. This could lead us to incorrectly conclude that most of the exosomal DNA is surface-exposed.

**Figure 4.**
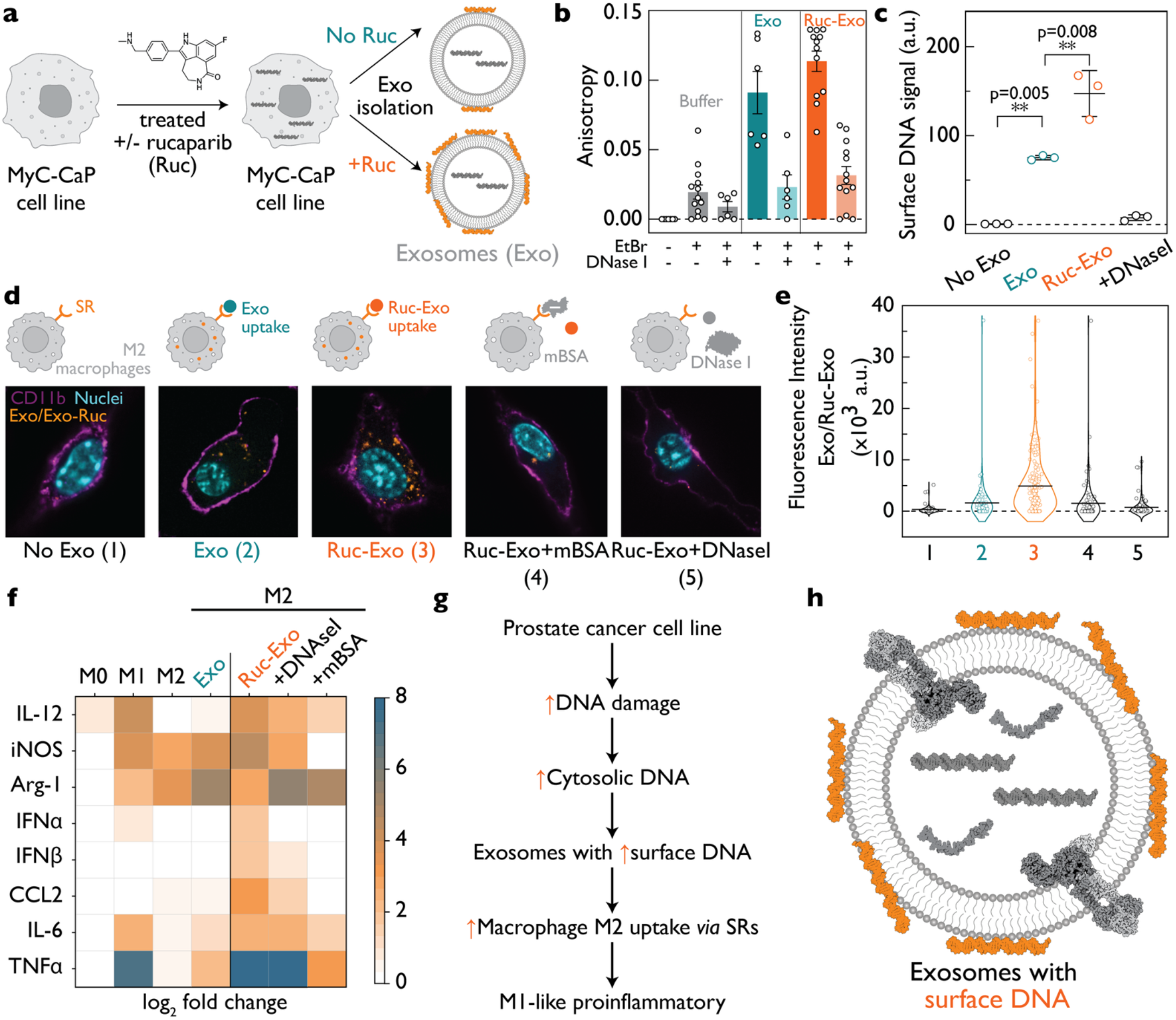
RUNA reveals presence of exosomal surface DNA and its role in exosome uptake and macrophage polarization. **(a)** Exosomes are isolated from Myc-CaP prostate cancer cells with (Ruc-Exo) or without (Exo) 0.5 µM rucaparib (Ruc) treatment (36 h). **(b)** Fluorescence anisotropy of EtBr labeled Exo and Ruc-Exo with or without DNase I treatment (n=64). **(c)** RUNA labeling with membrane-impermeant sulfo-Cy3 reveals surface DNA on Exo and Ruc-Exo (n=12). **(d)** Confocal images showing uptake of exosomes fluorescently labeled with DiO (yellow) by M2 macrophages. M2 macrophages are incubated with (1) medium (no Exo); (2) Exo (3) Ruc-Exo (4) Ruc-Exo and 30 equiv of mBSA. (5) Exo and DNase I. Scale bar: 5 µm. **(e)** Quantification of uptake in (1) - (5). Statistical significance of Exo (p=1.53 × 10^-5^) and Ruc-Exo (p=1.51 × 10^-7^) uptake compared to the no Exo. Statistical significance of Ruc-Exo with Ruc-Exo + DNase I (p = 2.25 × 10^-5^) and Ruc-Exo + mBSA (p = 2.91 × 10^-4^) (n=479 cells for all conditions). Uptake 10 are log_1+x_-transformed and analyzed by one-way ANOVA with Tukey’s HSD correction. Data are reported for single cells with group means. (**f**) M2 macrophages treated with Exo and Ruc-Exo exhibit M1-like cytokine and chemokine profiles, but not when pretreated with DNase I or mBSA. (**g**) Proposed model: Genotoxic stress induced by rucaparib elevates surface DNA on exosomes (**h**), promotes macrophage uptake via scavenger receptors and modulates macrophage immune response.

To distinguish between either possibility, we devised a covalent, membrane-delimited labeling strategy based on RUNA to pinpoint the localization of exosomal DNA. We used methyl isonitrile and norbornene aldehyde to conjugate the membrane-impermeant sulfo-Cy3 tetrazine to DNA (**Figure 4c, Figure S33**). RUNA was applied to equal concentrations of RNase A-treated exosomes from untreated (Exo) and rucaparib treated (Ruc-Exo) MyC-CaP cells. In all cases free unreacted dye was removed by stringent ultrafiltration, leaving only covalently attached dye molecules. Because the Cy3 dye is membrane-impermeable, and any lumenal DNA escapes modification, any Cy3 signal can be attributed solely to surface DNA. We found that Exo samples were substantially labeled with Cy3 and that DNase I treatment dramatically reduced the signal indicating that most exosomal DNA is surface-exposed. Surprisingly, this signal nearly doubled in Ruc-Exo samples and was completely lost when samples were pre-treated with DNase I. Nanoparticle tracking analysis (NTA) showed that exosome size distributions were unchanged after RUNA labeling and/or DNase I treatment, suggesting that these treatments do not measurably disrupt overall vesicle integrity (**Figure S34**). RUNA reveals that not only is most exosomal DNA surface-exposed, but its abundance on exosomes nearly doubles in response to the accumulation of DNA damage in cells.

Most solid tumors are macrophage-centric where tumor-associated macrophages (TAMs) account for nearly 50% of the total immune population (40, 41). Macrophages tend to adopt an M2-like anti-inflammatory phenotype within the tumor microenvironment which promotes tumor growth (42–44). To probe the functional relevance of surface DNA on exosomes, we tested their effect on M2-polarized macrophages. We labeled both kinds of exosomes with Dojindo Exosparkler (DiO), a lipophilic fluorescent dye that stains exosome membranes. We then pulsed M2 macrophages with equal amounts of Exo or Ruc-Exo and quantified exosome uptake by fluorescence microscopy (**Figure S35**-**S38)**. We found that uptake of Ruc-Exo was much higher than that of Exo by M2 macrophages (**Figure 4d**; **Table S2**). Scavenger receptors are highly expressed in macrophages and are known to bind and internalize anionic ligands such as duplex DNA (45–49). Competition experiments with excess maleylated BSA (mBSA) revealed that uptake was indeed predominantly *via* scavenger receptors. Furthermore, pre-treating Ruc-Exo with DNase I effectively abolished exosome uptake. This data establishes that surface DNA drives exosome uptake by engaging scavenger receptors on M2 macrophages.

To test whether exosome uptake had any functional impact on M2 macrophages, we profiled the cytokines they secreted before and after exosome uptake (**Figure 2f**). We observed that Ruc-Exo uptake led to M2 macrophages showing an increased expression of markers indicative of an activated M1 macrophage cell state. The secretion of type I interferon, IL-12, and TNF-α increased, as did the levels of iNOS and CCL2. Concomitantly, the levels of proteins such as arginase-1, which is abundant in anti-inflammatory M2 macrophages, decreased. The cytokine/chemokine profile for M2 macrophages treated with Ruc-Exo better resembles that of M1 macrophages, rather than canonical M2 macrophages or those that have taken up exosomes (Exo) from untreated cancer cells. Further, M2-macrophages that take up Ruc-Exo pretreated with DNase I or mBSA do not express most of these pro-inflammatory markers. Our results reveal that surface DNA on exosomes from drug-treated cancer cells shifts M2-polarized macrophages to a more M1-like state (**Figure 4h, Figure S38**).

## Discussion

RUNA is a new reversible, nucleobase-selective covalent chemistry that can discriminate nucleic acid populations, irrespective of their sequence, if they are separated by a charged biological membrane. Unlike classical carbodiimide reagents such as EDC or CMC, which irreversibly modify uridines and guanosines and eventually disrupt structure (4), RUNA exhibits a nucleobase preference for U and T in nucleic acids. Any apparent sequence preference would likely be governed by accessibility (2, 50). Unlike acylation-based methods that target only RNA (1), RUNA can be applied to both RNA and DNA. Importantly, one can reverse this modification and recover native RNA and DNA without loss of integrity, a feature that was previously limited to enzymatic or photocleavable systems.

A distinctive feature of RUNA is its spatial selectivity, which arises from its modular nature. It uses two simple and commercially available modules: aldehydes and isonitriles. By using a membrane-impermeant or a membrane permeant aldehyde, RUNA can chemically discriminate nucleic acid pools that are membrane-encapsulated or not. The advantage of using RUNA rather than a pair of membrane-permeable or -impermeable dyes is that RUNA decouples the fluorescence readout from membrane permeability. It can use same fluorophore for a readout because the aldehyde component determines membrane permeability. This bypasses photophysical differences that would result upon chemically modifying a dye to create membrane permeable and impermeable versions.

By applying this chemistry to exosomes, we resolved the previously ill-defined localization of DNA in exosomes, which has limited our understanding of exosome function (27, 31). Though prior studies have detected DNA in exosomes (27, 32, 51), it was not known whether it was encapsulated or exposed (27). This has been technically challenging to establish thus far because of the low abundance of exosomal DNA. Approaches such as qPCR, antibody-based detection, or UV spectroscopy are either potentially biased by differences in DNA accessibility, chemistry or sequence, or lack sufficient sensitivity. RUNA unambiguously shows that a significant fraction of exosomal DNA is surface-exposed and that it increases when cells are under genotoxic stress. This observation links DNA repair with exosome composition suggesting a bridge between genome integrity and exosome-mediated cell-cell communication.

Our findings point to a new role for exosomal DNA. Exosomal DNA is currently considered a passive cargo. However, we observe that exosomes leverage this DNA component for cell entry by engaging scavenger receptors. Further, after uptake, changes in the abundance or composition of exosomal DNA, or both, shift M2 macrophages toward an M1-like state. This suggests that tumor-derived exosomes can encode immunomodulatory signals in their nucleic acid content. Our observation may partly explain the immune-modulating effects of PARP inhibitors observed *in vivo* (52), where their activity extends beyond DNA damage repair to include modulation of the tumor microenvironment.

Several open questions remain regarding exosomal surface DNA. Studies across additional cell lines, primary tissue-derived exosomes, and macrophage models will be needed to determine whether surface DNA is a general feature of exosomes. It is currently unclear whether surface DNA represents a regulated biological signal or is a byproduct of cell damage. Fundamental properties such as its length, sequence composition, 11 modification profiles, whether single-stranded, double stranded or histone-complexed, and how each of these features influence immune recognition, remain to be defined. The biogenesis of surface-adhered DNA and its mode of attachment to exosomes are open questions.

RUNA labeling opens multiple new avenues of investigation. Chemically, it offers a versatile scaffold for temporal and spatial labeling of nucleic acids in living systems, potentially enabling pulse–chase imaging or reversible installation of functional tags. Biologically, it provides a platform to interrogate surface nucleic acids associated with biological membranes such as glycoRNAs, surface DNA on extracellular vesicles, and exosomes (24, 25, 29). This class of molecules is being increasingly implicated in cell–cell and immune communication, indicating that their chemical accessibility is a key determinant of their biological functionality (23, 28, 30). Future applications may include mapping RNA and DNA accessibility across organelle membranes, viruses, or synthetic protocells, and coupling reversible labeling with sequencing.

## MATERIALS AND METHODS

### RUNA Reaction

RUNA labeling reactions were performed in 20 µL aqueous solutions containing 1 µM Cy3-labeled RNA, 0-200 mM HEPES at the indicated pH, 10-200 mM isonitrile, and 10-200 mM aldehyde. Reactions were incubated at room temperature (18 °C) unless noted otherwise.

### Denaturing Electrophoretic Mobility Shift Assay

Reaction products were analyzed by 20 % (v/v) urea PAGE. Samples were mixed with formamide/EDTA loading buffer, heated, and resolved at constant power. Fluorescent bands were imaged using a Typhoon scanner, and band intensities were quantified using ImageQuant TL.

### ^1^H and ^31^P NMR Analysis

RUNA reactions for NMR were carried out in 500 µL solutions containing 25 mM nucleotide or nucleoside, 200 mM HEPES, 200 mM methyl isonitrile, 200 mM aldehyde, and 10 % D_2_O. Control reactions lacking individual components were prepared in parallel. Spectra were collected on a 400 MHz instrument using water suppression and analyzed in MestReNova.

### Phospholipid Vesicle Preparation for Extravesicular RNA Labeling

Phospholipid vesicles were prepared from POPC by thin-film rehydration in 200 mM HEPES (pH 8.0) containing 3′-ATTO647-labeled 12-nt RNA. Vesicles were sonicated and purified by size-exclusion chromatography on Sepharose 4B. Fractions were monitored by fluorescence to distinguish vesicle-encapsulated RNA from free dye. Vesicle size distributions and encapsulation efficiency were determined by dynamic light scattering.

### Cell Lines and Primary Cells

MyC-CaP prostate cancer cells were cultured in DMEM supplemented with 10 % FBS, 1 % penicillin– streptomycin, 2 % L-glutamine, and 0.2 % Plasmocin. Bone-marrow–derived macrophages (BMDMs) were generated from C57Bl/6J mice and differentiated for 5–7 d in DMEM containing 10 % FBS and 50 ng/mL M-CSF. All cell lines tested negative for mycoplasma.

### Exosome Assays and Rucaparib Treatment

MyC-CaP cells were treated with 500 nM rucaparib for 36 h before exosome collection. BMDMs were polarized to an M2-like state using 20 ng/mL IL-4 for 24 h and incubated with isolated exosomes for cytokine analysis, flow cytometry, or microscopy.

Conditioned media were sequentially centrifuged at 300 × g, 2,000 × g, and 10,000 × g, followed by ultracentrifugation at 100,000 × g for 70 min. Pelleted exosomes were washed once in PBS and quantified by nanoparticle tracking analysis. Size distributions were confirmed by dynamic light scattering.

### RUNA Labeling of Exosome Surface DNA

Exosomes (10^7^ particles) in 20 mM HEPES (pH 7.4) were incubated with 100 mM methyl isonitrile and 100 mM norbornene aldehyde at 4 °C for 12 h. Excess reagents were removed by centrifugal filtration (100 kDa MWCO). Tetrazine–sulfo-Cy3 was added to a final concentration of 5 mM for 2 h, followed by additional filtration. Fluorescence was measured on a Cary Eclipse fluorescence spectrometer (Agilent).

### Exosome Internalization Assays

Equal particle numbers of exosomes from control and rucaparib-treated cells were labeled using the Dojindo ExoSparkler membrane dye following the manufacturer’s instructions, with an additional ultrafiltration step to remove unbound dye. M2 macrophages were plated on glass-bottom dishes and incubated with labeled exosomes for 40 min. Cells were washed, fixed, and stained with anti-CD11b and Hoechst. Images were collected using a SoRa spinning-disk confocal microscope and analyzed in ImageJ. For inhibition experiments, macrophages were pre-treated with maleylated BSA or exosomes were pre-treated with DNase I.

### Intracellular Staining for Arginase-1 and iNOS / Cytokine, Chemokine Detection from Cell Supernatants

Macrophages were fixed and permeabilized using the BD Cytofix/Cytoperm protocol and stained with PE-conjugated anti-Arginase-1 and APC-conjugated anti-iNOS. Cells were analyzed on an LSR Fortessa cytometer, and data were processed in FlowJo. Cell supernatants were collected 24 h after rucaparib treatment (along with the controls) and used to detect analytes such as cytokines and chemokines using the protocol for LEGENDplex™ Mouse Anti-Virus Response Panel (BioLegend).

## ACKNOWLEDGMENTS

J.W.S. is an Investigator of the Howard Hughes Medical Institute. F.B. acknowledges funding from the European Molecular Biology Organization through a Long-Term Fellowship (ALTF 106-2023). Y.K. acknowledges NIH grants DP1GM149751, 1R01NS112139-01A1, 1R01GM147197-01, HFSP grant no: RGP0032/2022 and the Bimla Rani Parkinson’s Disease Research Fund. We are grateful to the Szostak lab members for their comments and insights on the manuscript.

## REFERENCES

1. R. C. Spitale, et al., Structural imprints in vivo decode RNA regulatory mechanisms. Nature 519, 486–490 (2015).

2. T. Wu, R. Lyu, Q. You, C. He, Kethoxal-assisted single-stranded DNA sequencing captures global transcription dynamics and enhancer activity in situ. Nat. Methods 17, 515–523 (2020).

3. N. Klöcker, F. P. Weissenboeck, A. Rentmeister, Covalent labeling of nucleic acids. Chem. Soc. Rev. 49, 8749–8773 (2020).

4. P. Y. Wang, A. N. Sexton, W. J. Culligan, M. D. Simon, Carbodiimide reagents for the chemical probing of RNA structure in cells. RNA 25, 135–146 (2019).

5. S. Nainar, et al., Temporal Labeling of Nascent RNA Using Photoclick Chemistry in Live Cells. J. Am. Chem. Soc. 139, 8090–8093 (2017).

6. Y. Motorin, et al., Expanding the chemical scope of RNA:methyltransferases to site-specific alkynylation of RNA for click labeling. Nucleic Acids Res. 39, 1943–1952 (2011).

7. L. Büttner, F. Javadi-Zarnaghi, C. Höbartner, Site-Specific Labeling of RNA at Internal Ribose Hydroxyl Groups: Terbium-Assisted Deoxyribozymes at Work. J. Am. Chem. Soc. 136, 8131–8137 (2014).

8. S. Croce, S. Serdjukow, T. Carell, T. Frischmuth, Chemoenzymatic Preparation of Functional Click-Labeled Messenger RNA. ChemBioChem 21, 1641–1646 (2020).

9. P. Wang, C. Ye, M. Zhao, B. Jiang, C. He, Small-molecule-catalysed deamination enables transcriptome-wide profiling of N6-methyladenosine in RNA. Nat. Chem. (2025). 10.1038/s41557-025-01801-3.

10. S. Nainar, et al., An optimized chemical-genetic method for cell-specific metabolic labeling of RNA. Nat. Methods 17, 311–318 (2020).

11. M. Rabani, et al., Metabolic labeling of RNA uncovers principles of RNA production and degradation dynamics in mammalian cells. Nat. Biotechnol. 29, 436–442 (2011).

12. C. Y. Jao, A. Salic, Exploring RNA transcription and turnover in vivo by using click chemistry. Proc. Natl. Acad. Sci. 105, 15779–15784 (2008).

13. K. Burger, et al., 4-thiouridine inhibits rRNA synthesis and causes a nucleolar stress response. RNA Biol. 10, 1623–1630 (2013).

14. F. Erhard, et al., Time-resolved single-cell RNA-seq using metabolic RNA labelling. Nat. Rev. Methods Primer 2, 77 (2022).

15. G. Vilkaitis, V. Masevičius, E. Kriukienė, S. Klimašauskas, Chemical Expansion of the Methyltransferase Reaction: Tools for DNA Labeling and Epigenome Analysis. Acc. Chem. Res. 56, 3188–3197 (2023).

16. G. M. Hanz, B. Jung, A. Giesbertz, M. Juhasz, E. Weinhold, Sequence-specific Labeling of Nucleic Acids and Proteins with Methyltransferases and Cofactor Analogues. J. Vis. Exp. 52014 (2014). 10.3791/52014.

17. J. Deen, et al., Methyltransferase-Directed Labeling of Biomolecules and its Applications. Angew. Chem. Int. Ed. 56, 5182–5200 (2017).

18. K. E. Deigan, T. W. Li, D. H. Mathews, K. M. Weeks, Accurate SHAPE-directed RNA structure determination. Proc. Natl. Acad. Sci. 106, 97–102 (2009).

19. R. Shapiro, J. Hachmann, The Reaction of Guanine Derivatives with 1,2-Dicarbonyl Compounds*. Biochemistry 5, 2799–2807 (1966).

20. M. Litt, V. Hancock, Kethoxal—A Potentially Useful Reagent for the Determination of Nucleotide Sequences in Single-Stranded Regions of Transfer Ribonucleic Acid. Biochemistry 6, 1848–1854 (1967).

21. R. Lyu, et al., KAS-seq: genome-wide sequencing of single-stranded DNA by N3-kethoxal–assisted labeling. Nat. Protoc. 17, 402–420 (2022).

22. P. T. Gilham, An Addition Reaction Specific for Uridine and Guanosine Nucleotides and its Application to the Modification of Ribonuclease Action. J. Am. Chem. Soc. 84, 687–688 (1962).

23. D. Jachertz, P. Anker, P. A. Maurice, M. Stroun, Information carried by the DNA released by antigen-stimulated lymphocytes. Immunology 37, 753–763 (1979).

24. M. Stroun, et al., Presence of RNA in the nucleoprotein complex spontaneously released by human lymphocytes and frog auricles in culture. Cancer Res. 38, 3546–3554 (1978).

25. S. A. Benner, Extracellular ‘communicator RNA.’ FEBS Lett. 233, 225–228 (1988).

26. L. Kageler, J. Perr, R. A. Flynn, Tools to investigate the cell surface: Proximity as a central concept in glycoRNA biology. Cell Chem. Biol. 31, 1132–1144 (2024).

27. T. Tsering, A. Nadeau, T. Wu, K. Dickinson, J. V. Burnier, Extracellular vesicle-associated DNA: ten years since its discovery in human blood. Cell Death Dis. 15, 668 (2024).

28. A. R. Thierry, S. El Messaoudi, P. B. Gahan, P. Anker, M. Stroun, Origins, structures, and functions of circulating DNA in oncology. Cancer Metastasis Rev. 35, 347–376 (2016).

29. R. A. Flynn, et al., Small RNAs are modified with N-glycans and displayed on the surface of living cells. Cell 184, 3109-3124.e22 (2021).

30. L. Murphy, et al., Platelets sequester extracellular DNA, capturing tumor-derived and free fetal DNA. Science 389, eadp3971 (2025).

31. A. Yokoi, et al., Mechanisms of nuclear content loading to exosomes. Sci. Adv. 5, eaax8849 (2019).

32. E. Lázaro-Ibáñez, et al., DNA analysis of low- and high-density fractions defines heterogeneous subpopulations of small extracellular vesicles based on their DNA cargo and topology. J. Extracell. Vesicles 8, 1656993 (2019).

33. A. Mariani, D. A. Russell, T. Javelle, J. D. Sutherland, A Light-Releasable Potentially Prebiotic Nucleotide Activating Agent. J. Am. Chem. Soc. 140, 8657–8661 (2018).

34. J. Zhang, et al., Efficient Assembly of Functional RNA by in Situ Phosphate Activation and Loop-Closing Ligation. J. Am. Chem. Soc. 147, 39212–39222 (2025).

35. F. Bošković, U. F. Keyser, Nanopore microscope identifies RNA isoforms with structural colours. Nat. Chem. 14, 1258–1264 (2022).

36. R. Kalluri, V. S. LeBleu, The biology, function, and biomedical applications of exosomes. Science 367, eaau6977 (2020).

37. E. Wahlberg, et al., Family-wide chemical profiling and structural analysis of PARP and tankyrase inhibitors. Nat. Biotechnol. 30, 283–288 (2012).

38. C. Kim, X.-D. Wang, Y. Yu, PARP1 inhibitors trigger innate immunity via PARP1 trapping-induced DNA damage response. eLife 9, e60637 (2020).

39. P. D. Gupta, et al., PARP and PI3K inhibitor combination therapy eradicates c-MYC-driven murine prostate cancers via cGAS/STING pathway activation within tumor-associated macrophages. (2020).

40. L. J. Dooling, et al., Clustered macrophages cooperate to eliminate tumors via coordinated intrudopodia. Proc. Natl. Acad. Sci. 122, e2425452122 (2025).

41. S. Yang, S. Wei, F. Wei, Extracellular vesicles mediated gastric cancer immune response: tumor cell death or immune escape? Cell Death Dis. 15, 377 (2024).

42. M. Zhang, et al., Alpha fetoprotein promotes polarization of macrophages towards M2-like phenotype and inhibits macrophages to phagocytize hepatoma cells. Front. Immunol. 14, 1081572 (2023).

43. K. Hamidzadeh, A. T. Belew, N. M. El-Sayed, D. M. Mosser, The transition of M-CSF–derived human macrophages to a growth-promoting phenotype. Blood Adv. 4, 5460–5472 (2020).

44. M. L. Lundahl, et al., Macrophage innate training induced by IL-4 and IL-13 activation enhances OXPHOS driven anti-mycobacterial responses. eLife 11, e74690 (2022).

45. B. Suresh, et al., Tubular lysosomes harbor active ion gradients and poise macrophages for phagocytosis. Proc. Natl. Acad. Sci. 118, e2113174118 (2021).

46. S. Surana, J. M. Bhat, S. P. Koushika, Y. Krishnan, An autonomous DNA nanomachine maps spatiotemporal pH changes in a multicellular living organism. Nat. Commun. 2, 340 (2011).

47. M. S. Jani, J. Zou, A. T. Veetil, Y. Krishnan, A DNA-based fluorescent probe maps NOS3 activity with subcellular spatial resolution. Nat. Chem. Biol. 16, 660–666 (2020).

48. C. Cui, et al., A lysosome-targeted DNA nanodevice selectively targets macrophages to attenuate tumours. Nat. Nanotechnol. 16, 1394–1402 (2021).

49. J. Zou, et al., A DNA nanodevice for mapping sodium at single-organelle resolution. Nat. Biotechnol. 42, 1075–1083 (2024).

50. L. Fang, L. Xiao, Y. W. Jun, Y. Onishi, E. T. Kool, Reversible 2′-OH acylation enhances RNA stability. Nat. Chem. 15, 1296–1305 (2023).

51. G. Shelke, S. C. Jang, Y. Yin, C. Lässer, J. Lötvall, Human mast cells release extracellular vesicle-associated DNA. Matters Zür. (2016). 10.19185/matters.201602000034.

52. A. K. Mehta, et al., Targeting immunosuppressive macrophages overcomes PARP inhibitor resistance in BRCA1-associated triple-negative breast cancer. Nat. Cancer 2, 66–82 (2020).

