## Supplementary Materials for "A nucleic acid labeling chemistry reveals surface DNA on exosomes"

**This PDF file includes**

Supplementary Materials and Methods
Supplementary References
Figures S1 to S38
Tables S1 to S3

**Supplementary Materials and Methods**

***Materials***

All chemicals were purchased from Ambeed, Combi-Blocks, or Sigma Aldrich unless otherwise noted. Specifically, methyl isonitrile (MeNC) was purchased from Sigma-Aldrich (ENAH58168694).

***Solid-State RNA Synthesis***

RNA oligonucleotides were synthesized on a K&A H6 oligosynthesizer using standard A, C, G, and U phosphoramidites (ChemGenes) and an N3-methyl-U phosphoramidite (Glen Research). A standard synthesis protocol was followed for RNA deprotection and Cyanine-3-containing oligonucleotide synthesis, respectively (1, 2). The RNA oligonucleotides were cleaved from the solid support by flushing fifteen times in both directions with 1.2 mL of freshly prepared ammonium hydroxide:methylamine (1:1, v/v) solution (AMA). The liquid was retained in the syringes for 15 min at room temperature (20 °C), protected from light, and then transferred to 2-mL screwcap microtubes. The column was further flushed with 1 mL of AMA, and the eluate was collected in a separate 2-mL microtube. The pink color of Cyanine-3 served as a visual indicator of complete cleavage. The microtubes were incubated for 15 min at 65 °C to remove base-protecting groups, cooled on ice for 3 min, centrifuged briefly, fitted with 23-gauge vented caps, and dried in a SpeedVac for 1.5 h. The remaining liquid was frozen in liquid nitrogen and lyophilized overnight.

To remove the 2′-O-TBDMS protecting groups, the dried oligonucleotides were dissolved in 100 µL DMSO, followed by the addition of 125 µL THF–TEA solution. The mixture was vortexed, briefly centrifuged, and incubated for 2.5 h at 65 °C, protected from light. The samples were then cooled on ice for 3 min and centrifuged. A 25 µL aliquot of 5 M ammonium acetate and 1 mL of ice-cold isopropanol were added, followed by vortexing and incubation on dry ice for 15 min. The samples were centrifuged at 16,000 × g for 15 min at 4 °C, and the supernatant was discarded, leaving a visible pink pellet. The pellet was washed with 1 mL of ice-cold 80 % (v/v) ethanol, centrifuged again at 16,000 × g for 15 min at 4 °C, and the supernatant was discarded. The RNA pellets were dissolved in 100 µL of 98 % (v/v) formamide containing 5 mM EDTA and then subjected to preparative gel purification.

***Preparative Gel Purification of Oligonucleotides***

The dried oligonucleotide pellets were dissolved and purified by denaturing polyacrylamide gel electrophoresis on a 20 % (v/v) polyacrylamide–urea gel. The gel solution was prepared by combining 64 mL of urea concentrate, 24 mL of urea diluent, and 10 mL of 10× TBE buffer (all from National Diagnostics). N,N,N′,N′-tetramethylethylenediamine (TEMED) (70 µL) was added and mixed thoroughly, followed by the addition of 250 µL of 10 % (w/v) ammonium persulfate. The solution was mixed again and poured into a glass casting cassette (20 × 20 cm) fitted with 1.5 mm spacers and a one-well comb. The gel was allowed to polymerize for 30 min, then pre-run for 10 min at 8 W in 1× TBE buffer.

The sample (100 µL of the oligonucleotide in formamide–EDTA solution) was carefully loaded at the bottom of the well. Electrophoresis was carried out for 10 min at 8 W, followed by separation at 25 W until the Cyanine-3–labeled oligonucleotide band reached the midpoint of the gel. The gel cassette was then disassembled, and the fluorescent band corresponding to the full-length oligonucleotide was excised and transferred to a 50 mL conical tube. The gel slice was fragmented by shaking, vortexing, or crushing. Extraction was performed by adding 20–25 mL of 5 mM ammonium acetate containing 5 mM EDTA, protecting the tube from light with aluminum foil, and rotating vertically overnight at 10-15 rpm.

***Oligonucleotide Desalting***

Sep-Pak C18 cartridges (Waters) were used for oligonucleotide desalting following gel purification. Each cartridge was sequentially washed with 20 mL of acetonitrile and 20 mL of 100 mM triethylammonium acetate (TEAA) buffer. The filtered (5 µm) solution containing the purified oligonucleotide was then loaded dropwise onto the cartridge and pushed through. The flow-through was collected in the same 50 mL conical tube, and the loading step was repeated once to ensure complete binding to the cartridge. The cartridge was subsequently washed twice with 10 mL of 10 mM TEAA, and the oligonucleotide was eluted into a 2 mL microcentrifuge tube using 2 mL of a 1:1 (v/v) mixture of methanol and acetonitrile.

The eluate was dried in a SpeedVac concentrator for 2 h, frozen in liquid nitrogen, and lyophilized overnight. The resulting oligonucleotide was dissolved in 100 µL of nuclease-free water and stored at -80 °C until use. Oligonucleotide concentration was determined by measuring a 1:10 dilution on a NanoDrop UV spectrophotometer. A working stock solution (10 µM) was prepared and stored at -20 °C.

***RUNA Reaction***

RUNA labeling of synthetic RNA oligonucleotides was performed in a 20 µL aqueous reaction containing 1 µM Cyanine-3 (Cy3)–labeled RNA, 200 mM 4-(2-hydroxyethyl)-1-piperazineethanesulfonic acid (HEPES) buffer at the indicated pH, 200 mM MeNC, and 200 mM aldehyde. Reactions were carried out at room temperature (18 °C).

***Denaturing Polyacrylamide Gel Shift Assay***

Denaturing polyacrylamide gel electrophoresis (PAGE) was performed using a 20 % (v/v) urea–polyacrylamide gel cast with 1 mm spacers and a comb. One microliter of the RUNA reaction mixture was combined with 9 µL of 98 % (v/v) formamide containing 5 mM ethylenediaminetetraacetic acid (EDTA). A 1–1.5 µL aliquot of this mixture was loaded onto the gel, which had been pre-run for 30 min at 10 W, and electrophoresis was carried out for 1.5–2.5 h at 25 W.

Following electrophoresis, the gel (in glass plates) was imaged using a Typhoon fluorescence scanner with the appropriate detection channels (Cy2 for ATTO488, Cy3 for Cyanine-3, and Cy5 for ATTO647). Band intensities were quantified using ImageQuant TL software (Cytiva).

***$^{1}$H and $^{31}$P Nuclear Magnetic Resonance (NMR)***

RUNA reactions for NMR analysis were prepared in 500 µL aqueous solution containing 25 mM nucleotide or nucleoside, 200 mM HEPES buffer at the indicated pH, 200 mM isonitrile, 200 mM aldehyde, and 10 % (v/v) deuterium oxide ($D_2O$). Reactions were incubated at 18 °C. All NMR spectra were acquired on a 400 MHz spectrometer in water suppression mode. For each nucleotide or nucleoside, six samples were analyzed: (1) a complete reaction mixture lacking the nucleotide or nucleoside, (2) a mixture lacking aldehyde, (3) a mixture lacking MeNC, (4) a mixture lacking both MeNC and aldehyde, and (5, 6) two reactions containing all components, one of which was heated at 95 °C for 15 min in a preheated water bath after the labeling reaction. All samples were analyzed by $^{1}$H and $^{31}$P NMR spectroscopy to determine the extent of nucleotide modification and the reversibility of the reaction upon heating. Spectra were processed and analyzed using MestReNova software, version 14.2.0 (Mestrelab Research).

***Phospholipid Vesicle Preparation***

1-Palmitoyl-2-oleoyl-sn-glycero-3-phosphocholine (16:0–18:1 PC; POPC) was obtained from Avanti Polar Lipids and dissolved in chloroform at a concentration of 25 mg/mL. POPC vesicles were prepared by the thin-film rehydration method (3). A 250 µL aliquot of the 25 mg/mL POPC solution was transferred to a glass tube and dried under a continuous stream of nitrogen in a fume hood to form a thin lipid film. The tube was vacuumed for at least 2 h to remove residual solvent. The dried lipid film was rehydrated with 100 µL of 200 mM HEPES buffer, pH 8.0, containing 30 µM of a 12-nucleotide RNA oligonucleotide labeled at the 3′ end with ATTO647. The resulting suspension was vortexed thoroughly and sonicated for 30 min to form small unilamellar vesicles.

***Size Exclusion Chromatography (SEC) Vesicle Purification***

A size exclusion column was prepared by manually loading 5 mL of Sepharose 4B (4) ethanol slurry into a plastic column. The column was allowed to drain and then washed twice with 10 mL of MilliQ water, followed by one wash with 15 mL of 200 mM HEPES buffer, pH 8.0. A 100 µL aliquot of the vesicle suspension was applied to the column and allowed to fully enter the gel bed. Elution was carried out with fresh HEPES buffer, and the effluent was collected as fractions of approximately five drops per well in a 96-well microplate.

After the elution of the dye-labeled fraction (typically within the first 48 wells), fluorescence was measured using a microplate reader. The excitation and emission wavelengths were optimized using an ATTO647-containing well before analyzing all fractions. The resulting chromatogram showed two distinct peaks: an early-eluting population corresponding to vesicles encapsulating ATTO647-labeled RNA (fractions 9–13) and a later-eluting population corresponding to free dye.

***Protocell Characterization by Dynamic Light Scattering (DLS)***

The size distribution of the protocells was analyzed by DLS with a Malvern Zetasizer instrument. Measurements were performed on both buffer and vesicle fractions to determine vesicle diameter.

***Membrane-Delimited RUNA Assay***

ATTO647-labeled RNA-containing vesicles (40 µL) were mixed with 1 µL of 30 µM ATTO488-labeled RNA. MeNC and aldehyde were then added stepwise in 10 mM increments to final concentrations of 200 mM each in a total reaction volume of 100 µL. Four parallel reactions were performed: (1) a control containing only 200 mM MeNC, and (2–4) reactions containing 200 mM MeNC with 200 mM acetaldehyde, norbornene aldehyde, or betaine aldehyde, respectively. The reactions were incubated for 6 h at 18 °C and subsequently purified by SEC using freshly prepared Sepharose 4B columns. Fractions containing ATTO647-labeled RNA (encapsulated vesicles) and ATTO488-labeled RNA (extravesicular RNA) were collected in a 96-well microplate and transferred to 1.5 mL microtubes.

To lyse vesicles, 35 µL of 20 % (v/v) Triton X-100 was added to each fraction, followed by 850 µL of chloroform. The mixtures were vortexed thoroughly, and the upper aqueous phases were transferred to new microcentrifuge tubes. Samples were concentrated in a SpeedVac for 30 min, frozen in liquid nitrogen, and lyophilized overnight. The dried material was resuspended in 10 µL of nuclease-free water, vortexed, and sonicated for 2 min. One microliter of each sample was mixed with 9 µL of 98 % (v/v) formamide containing 5 mM ethylenediaminetetraacetic acid (EDTA), and 1.5 µL of this mixture was analyzed by PAGE as described above.

***Cell Lines and Primary Cells***

The murine prostate cancer cell line Myc-CAP was obtained from the American Type Culture Collection (ATCC) and maintained in Dulbecco's Modified Eagle's Medium (DMEM) without phenol red, supplemented with 10 % fetal bovine serum (FBS), 1 % penicillin–streptomycin (P/S), 2 % L-glutamine, and 0.2 % Plasmocin (InvivoGen). Bone marrow–derived monocytes (BMDMs) were isolated from C57BL/6J mice and differentiated into macrophages in DMEM supplemented with 10 % FBS, 1 % P/S, 2 % L-glutamine, 0.2 % Plasmocin, and 50 ng/mL recombinant macrophage colony-stimulating factor (M-CSF) for 5–7 days. All cell lines were tested using the Universal Mycoplasma Detection Kit (ATCC) and confirmed to be mycoplasma-free.

***In Vitro Assays with Rucaparib and Exosomes***

Myc-CAP cells were treated with 500 nM Rucaparib for 36 h, and supernatants were collected for exosome isolation. Macrophages derived from BMDMs were polarized into M2-like cells by supplementation with 20 ng/mL recombinant murine interleukin-4 (IL-4) for 24 h. M2 macrophages were then co-incubated with $10^7$ exosome (Exo) particles isolated from control or Rucaparib-treated Myc-CAP cultures. After incubation, cell-free supernatants were collected for cytokine and chemokine analysis using the LEGENDplex™ Mouse Anti-Virus Response Panel (BioLegend), according to the manufacturer's instructions. Cells were used for intracellular staining of arginase-1 and inducible nitric oxide synthase (iNOS). For confocal microscopy, M2 macrophages were incubated with exosomes for 40 min, whereas for functional assays, incubation continued for 22–23 h.

***Exosome Isolation from Cancer Cell Culture Supernatant***

Myc-CAP cells were either left untreated or treated with 500 nM Rucaparib for 36 h. Cell culture supernatants were collected and centrifuged at 300 × g for 5 min at 4 °C to remove intact cells, followed by centrifugation at 2,000 × g for 10 min and 10,000 × g for 30 min at 4 °C to eliminate debris and larger vesicles. The cleared supernatant was then ultracentrifuged at 100,000 × g for 70 min at 4 °C using a Type 60 Ti rotor (38,000 rpm). The resulting exosome pellet was resuspended in 2 mL of phosphate-buffered saline (PBS), washed once by repeating the ultracentrifugation step, and resuspended in PBS for quantification.

Particle size distribution and concentration were determined using a NanoSight LM10 HS-BF instrument (NanoSight Ltd., UK) equipped with a 405 nm, 65 mW laser and EMCCD camera, based on nanoparticle tracking analysis (NTA). Exosomes were diluted 1:500 in particle-free PBS (pH 7.4) to achieve the optimal concentration range for NTA. The resulting exosomes exhibited a mean size of 100–200 nm. DLS analysis using a DynaPro Nanostar instrument (Wyatt Technologies) confirmed the size range. All measurements were performed in triplicate (n = 3), acquiring at least 5,000 events per sample. Equal numbers of exosomes from each condition were used for downstream experiments.

***Dot Blot Assay for Characterization of Exosomes***

Protein concentration for each group – exosome fraction, cell lysate was determined using PierceTM BCA Protein Assay Kit (ThermoFisher Scientific). Equal concentrations (by protein) of cell lysate and exosome fraction were spotted on nitrocellulose (NC) membrane and allowed to dry completely. The membranes underwent a blocking step with bovine serum albumin (BSA) in 5% TBST (Tris Buffered Saline with Tween-20) for 10-20 mins, followed by incubation with primary anti-mouse antibodies against ALIX, CD9, HSP70, β-actin at recommended dilutions (Cell Signaling Technology) for 1 hr at room temperature. After incubation with the primary antibody, the membranes were washed with 5% TBST, twice for 20-30 mins each time. The dot blot membrane was then incubated with HRP-conjugated anti-rabbit secondary antibody (Cell Signaling Technology) for 30 mins., followed by 2-3 additional wash steps with 5% TBST. Finally, the blots in 5% TBST were developed using chemiluminescent PierceTM ECL blotting substrate (ThermoFisher Scientific). All images were acquired on a BIO-RAD Imager and densitometry-based quantification was done using Image-Lab (BIO-RAD) software.

#### *RUNA Labeling of Exosome Surface DNA*

RNA-free exosomes ($10^7$ particles, normalized by NTA) purified in 20 mM HEPES buffer, pH 7.4, were incubated with MeNC and norbornene aldehyde, added incrementally over 15 min in equal 10 mM aliquots (final concentration 100 mM each). Reactions proceeded for 12 h at 4 °C. Excess reagents were removed by buffer exchange using a 4 mL Amicon® Ultra 100 kDa centrifugal filter (Millipore) with three washes in 4 mL of HEPES buffer until free reactant concentrations were below 0.01 mM. Labeled exosomes were then incubated with tetrazine–sulfo-Cy3 (5 mM; Lumiprobe) (5) for 2 h at 4 °C, followed by three additional washes with HEPES buffer using the same filtration procedure. Fluorescence intensity was measured using a Cary Eclipse fluorometer (Agilent Technologies) with excitation optimized at 498 nm.

#### *Labeling of Exosomes for Internalization Assay*

Equal numbers of exosomes from control and Rucaparib-treated cells were labeled using the ExoSparkler Exosome Membrane Labeling Kit (Dojindo, catalog no. EX03) according to the manufacturer's instructions. To remove unbound dye, samples were washed using Amicon® Ultra centrifugal filters (3 kDa molecular weight cutoff, Millipore) before use.

#### *Exosome Internalization Assay and Confocal Microscopy*

M2 macrophages were seeded on glass-bottom dishes at a density of $2 \times 10^3$ cells per well one day before the assay. Cells were incubated for 40 min with labeled exosomes from control or treated groups, washed with PBS, fixed with 4 % paraformaldehyde (PFA), and co-stained with anti-CD11b antibody and Hoechst nuclear stain. Samples were imaged using a SoRa subdiffraction Marianas spinning disk confocal microscope equipped with a 63× oil immersion objective (Intelligent Imaging Innovations). Images were processed and analyzed with ImageJ software.

For specific experimental conditions: (a) in maleylated bovine serum albumin (mBSA) inhibition assays, M2 macrophages were pretreated with 30 molar equivalents of mBSA for 30 min at 37 °C before exposure to labeled exosomes; (b) in DNase I degradation assays, exosomes were treated with DNase I (Thermo Fisher Scientific, RNase-free, 1 U/µL) following the manufacturer's instructions, then labeled using the ExoSparkler kit before incubation with macrophages.

#### *Arginase-1, iNOS, Cytokine and Chemokine Detection*

Macrophages were fixed and permeabilized using the BD Cytofix/Cytoperm protocol and stained with PE-conjugated anti-Arginase-1 and APC-conjugated anti-iNOS. Cells were analyzed on an LSR Fortessa cytometer, and data were processed in FlowJo. Cell supernatants were collected 24 hours after Rucaparib treatment (along with the controls) and used for detection of analytes such as cytokines and chemokines using the protocol for the LEGENDplex™ Mouse Anti-Virus Response Panel (BioLegend).

Briefly, this is a bead-based multiplex assay utilizing fluorescence–encoded beads suitable for simultaneous detection of multiple cytokines and chemokines, which can be analyzed using flow cytometry. The kits come with capture and detection antibodies as well as standards for the assay, which are serially diluted to generate a standard curve. The capture beads are coated with antibodies against target analytes and allow for simultaneous detection of multiple target proteins in cell supernatants, as well as reconstituted standards. Following incubation of samples and standards with capture beads for the recommended time, the beads are washed with 1× wash buffer (supplied with the kit) and further incubated with biotinylated detection antibodies. Next, secondary Streptavidin-phycoerythrin (SA-PE) is added to both standards and samples for the recommended incubation time. Both samples and standards are acquired as per recommended settings on the flow cytometer. Analysis of the results was performed using the proprietary software provided with the kit.

**Supplementary References**


1. The Glen Report. 21, 12–13 (2009).
2. The Glen Report. 25, 8–9 (2013).
3. C. Kirby, G. Gregoriadis, Dehydration-Rehydration Vesicles: A Simple Method for High Yield Drug Entrapment in Liposomes. *Nat Biotechnol* 2, 979–984 (1984).
4. T. Ruysschaert, *et al.*, Liposome retention in size exclusion chromatography. *BMC Biotechnol* 5, 11 (2005).
5. G. Devi, A. K. Hedger, R. J. Whitby, J. K. Watts, Double Click: Unexpected 1:2 Stoichiometry in a Norbornene–Tetrazine Reaction. *J. Org. Chem.* 88, 5341–5347 (2023).

**Fig.S1**

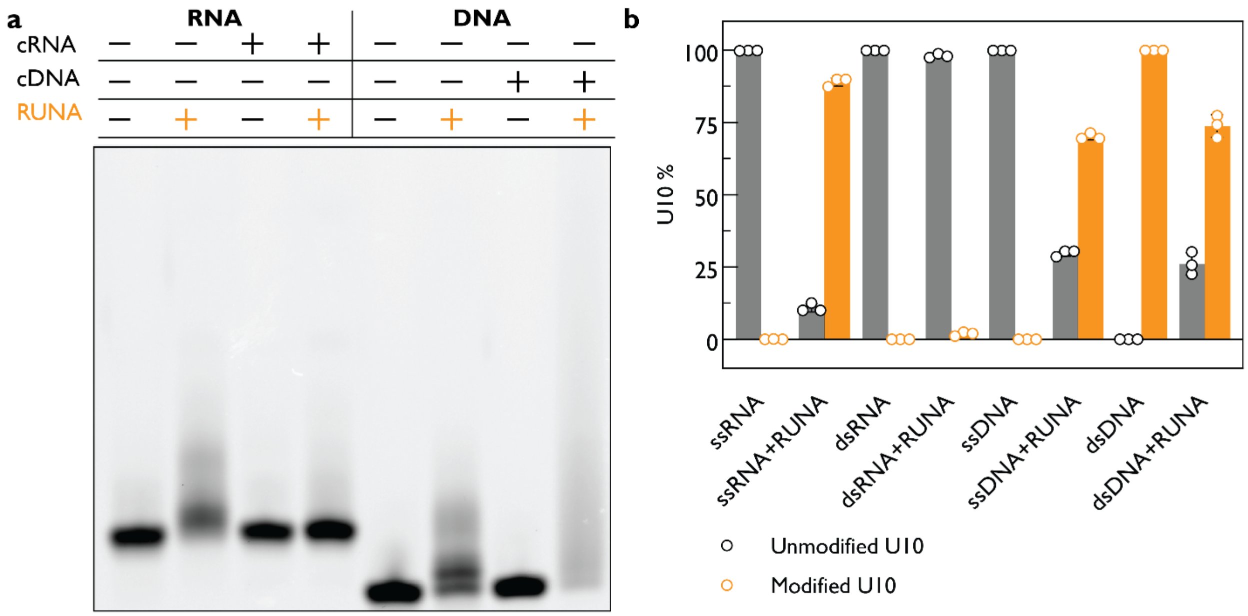


**Figure S1.** Single-stranded and double-stranded RNA and DNA 10-mers (1.5 µM; 3′-Cy3-labeled) reacted under RUNA conditions (100 mM MeNC, 100 mM 4-pentenal, 50 mM HEPES, pH 8.0, 12 h, 18 °C). Complementary RNA (cRNA) and complementary DNA (cDNA) strands (5 µM) were annealed before labeling in 10 mM $MgCl_2$ and 50 mM HEPES, pH 8.0, by heating for 3 min at 70 °C followed by slow cooling to 20 °C over 40 min. **(a)** Denaturing gel showing control and RUNA-treated samples; **(b)** corresponding quantification of fluorescence signal.

**Fig.S2**

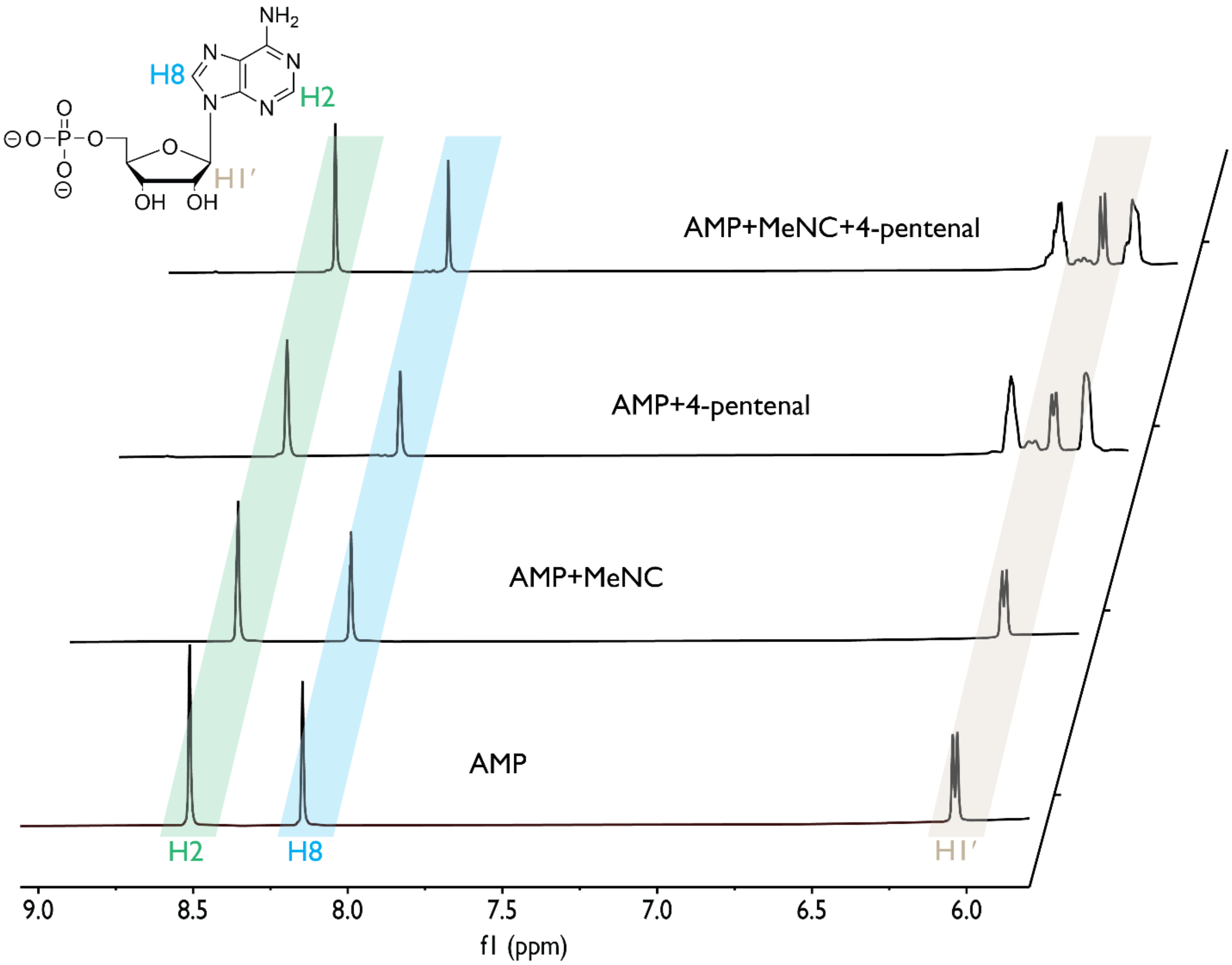


**Figure S2**. Adenosine 5′-monophosphate (AMP) $^{1}$H NMR spectra acquired in 200 mM HEPES, pH 8.0, in 9:1 (v/v) $H_2O$:$D_2O$ after 24 h. (1) 25 mM AMP; (2) 25 mM AMP, 100 mM MeNC; (3) 25 mM AMP, 100 mM 4-pentenal; (4) 25 mM AMP, 100 mM MeNC, 100 mM 4-pentenal. Newly observed peaks in (3) correspond to 4-pentenal.

**Fig.S3**

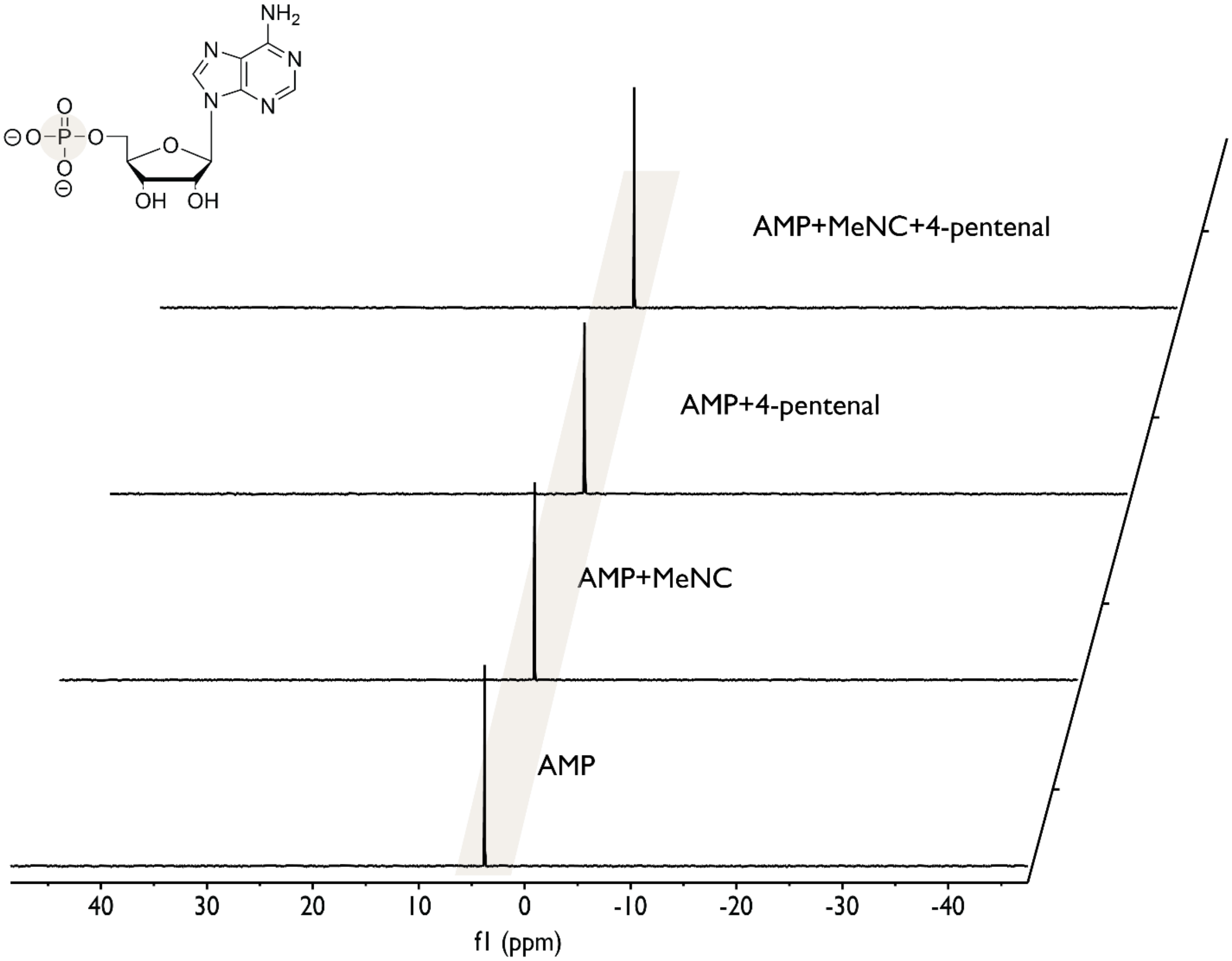


**Figure S3**. Adenosine 5′-monophosphate (AMP) $^{31}P$ NMR spectra acquired in 200 mM HEPES, pH 8.0, in 9:1 (v/v) $H_2O$:$D_2O$ after 24 h. (1) 25 mM AMP; (2) 25 mM AMP, 100 mM MeNC; (3) 25 mM AMP, 100 mM 4-pentenal; (4) 25 mM AMP, 100 mM MeNC, 100 mM 4-pentenal. No phosphate modification is detected.

**Fig.S4**

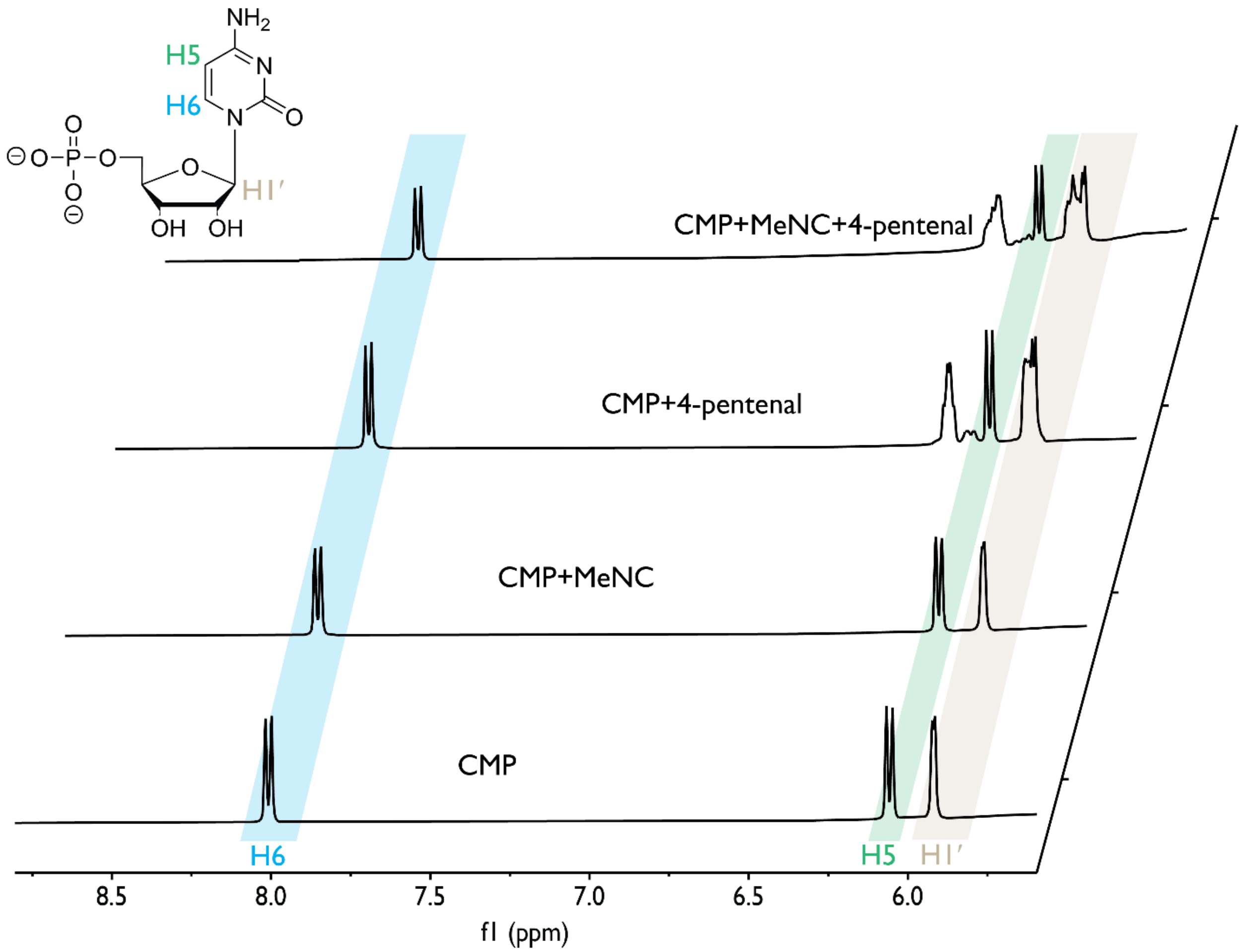


**Figure S4**. Cytidine 5′-monophosphate (CMP) [1]H NMR spectra acquired in 200 mM HEPES, pH 8.0, in 9:1 (v/v) $H_2O$:$D_2O$ after 24 h. (1) 25 mM CMP; (2) 25 mM CMP, 100 mM MeNC; (3) 25 mM CMP, 100 mM 4-pentenal; (4) 25 mM CMP, 100 mM MeNC, 100 mM 4-pentenal.

**Fig.S5**

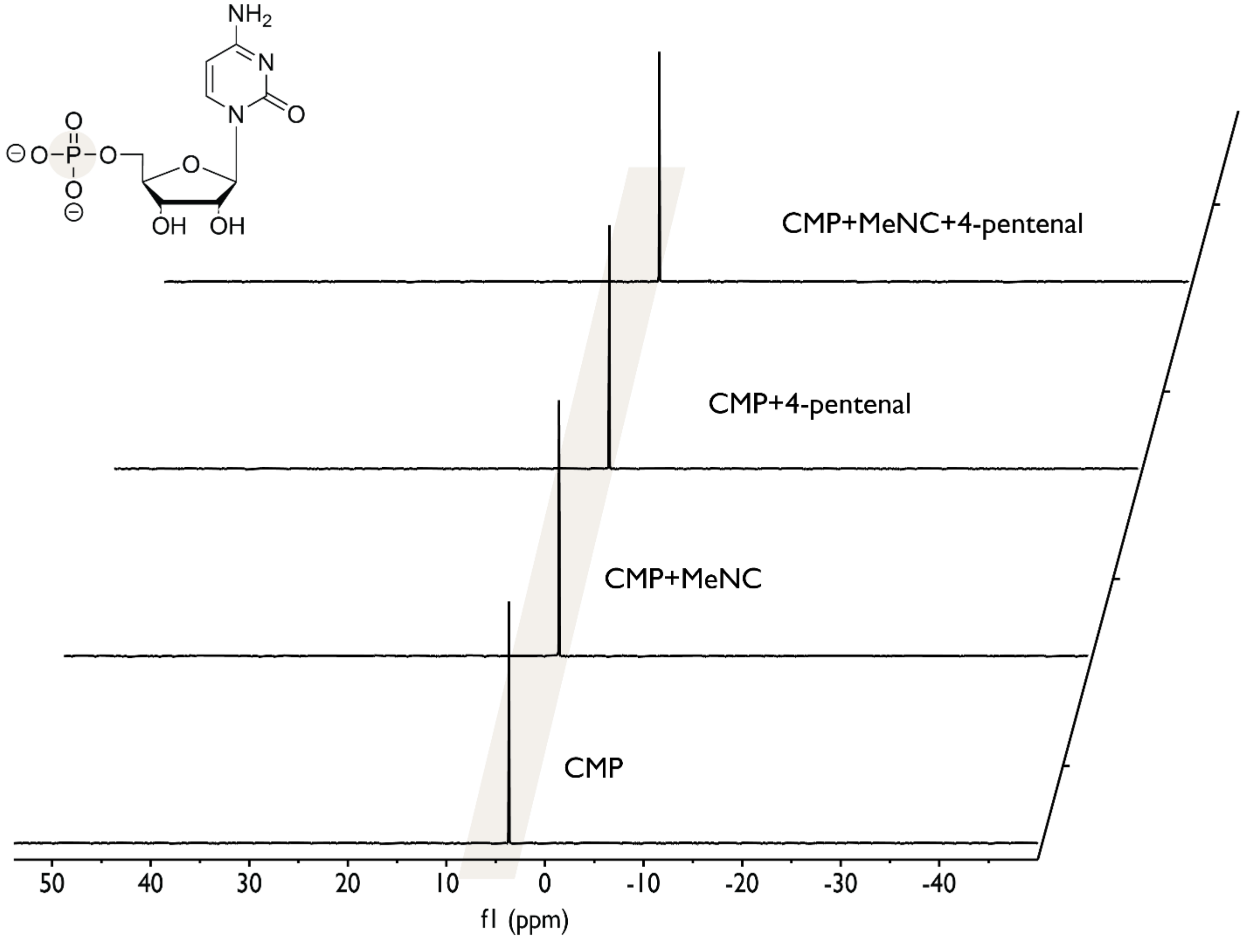


**Figure S5**. Cytidine 5′-monophosphate (CMP) $^{31}P$ NMR spectra acquired in 200 mM HEPES, pH 8.0, in 9:1 (v/v) $H_2O$:$D_2O$ after 24 h. (1) 25 mM CMP; (2) 25 mM CMP, 100 mM MeNC; (3) 25 mM CMP, 100 mM 4-pentenal; (4) 25 mM CMP, 100 mM MeNC, 100 mM 4-pentenal. No phosphate modification is detected.

**Fig.S6**

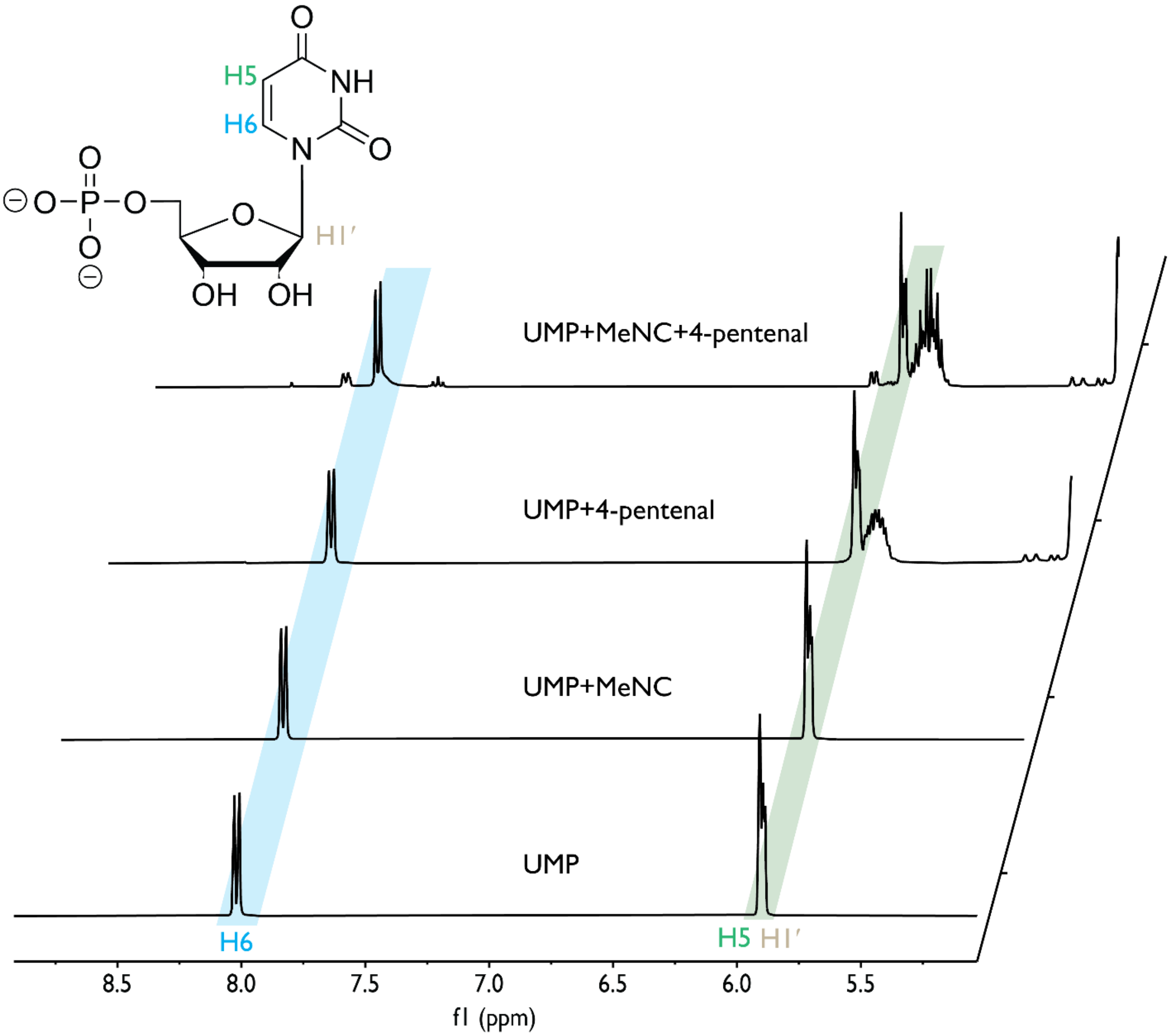


**Figure S6**. Uridine 5′-monophosphate (UMP) $^{1}$H NMR spectra acquired in 200 mM HEPES, pH 8.0, in 9:1 (v/v) $H_2O$:$D_2O$ after 24 h. (1) 25 mM UMP; (2) 25 mM UMP, 100 mM MeNC; (3) 25 mM UMP, 100 mM 4-pentenal; (4) 25 mM UMP, 100 mM MeNC, 100 mM 4-pentenal.

**Fig.S7**

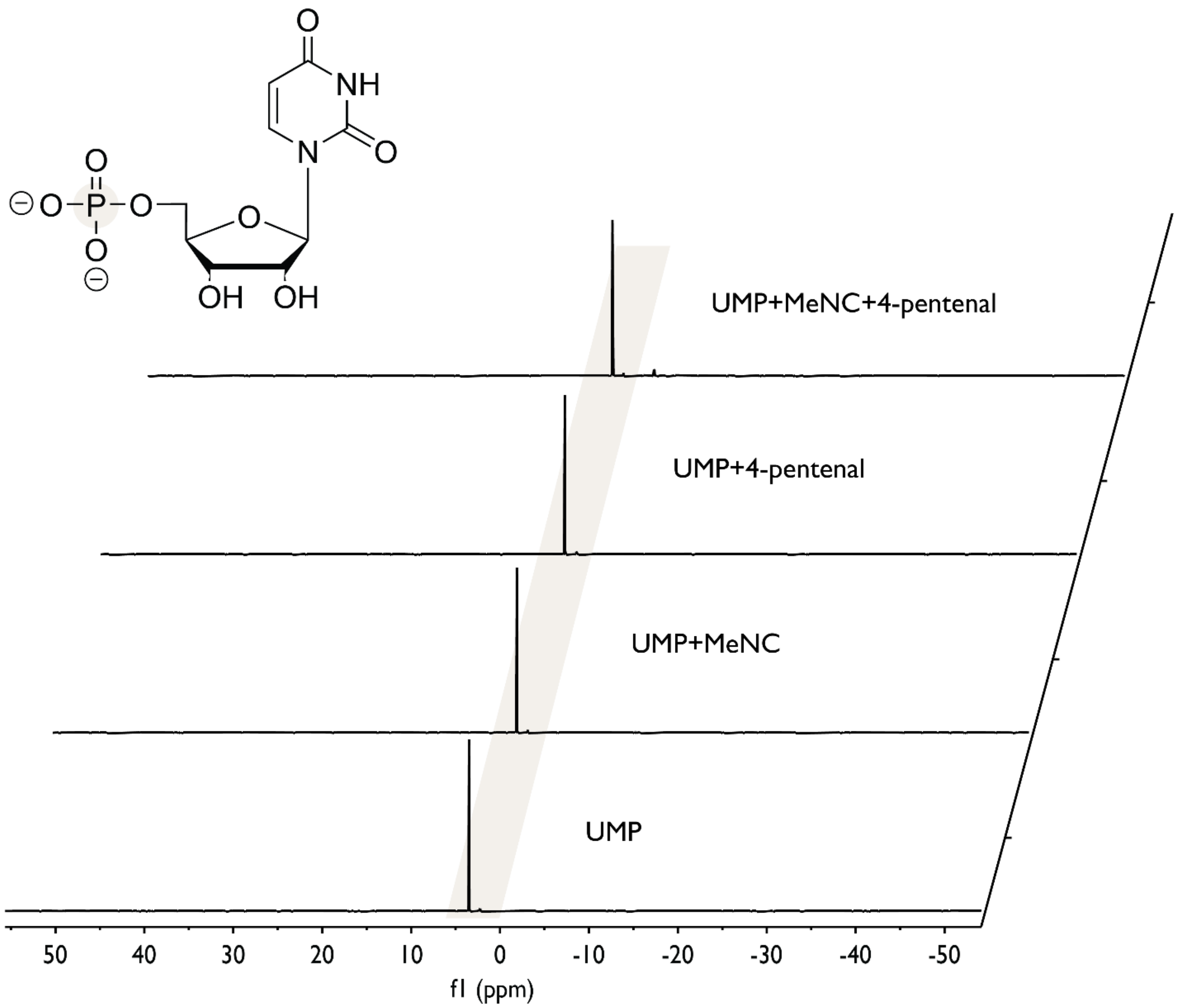


**Figure S7**. Uridine 5′-monophosphate (UMP) $^{31}P$ NMR spectra acquired in 200 mM HEPES, pH 8.0, in 9:1 (v/v) $H_2O$:$D_2O$ after 24 h. (1) 25 mM UMP; (2) 25 mM UMP, 100 mM MeNC; (3) 25 mM UMP, 100 mM 4-pentenal; (4) 25 mM UMP, 100 mM MeNC, 100 mM 4-pentenal.

**Fig.S8**

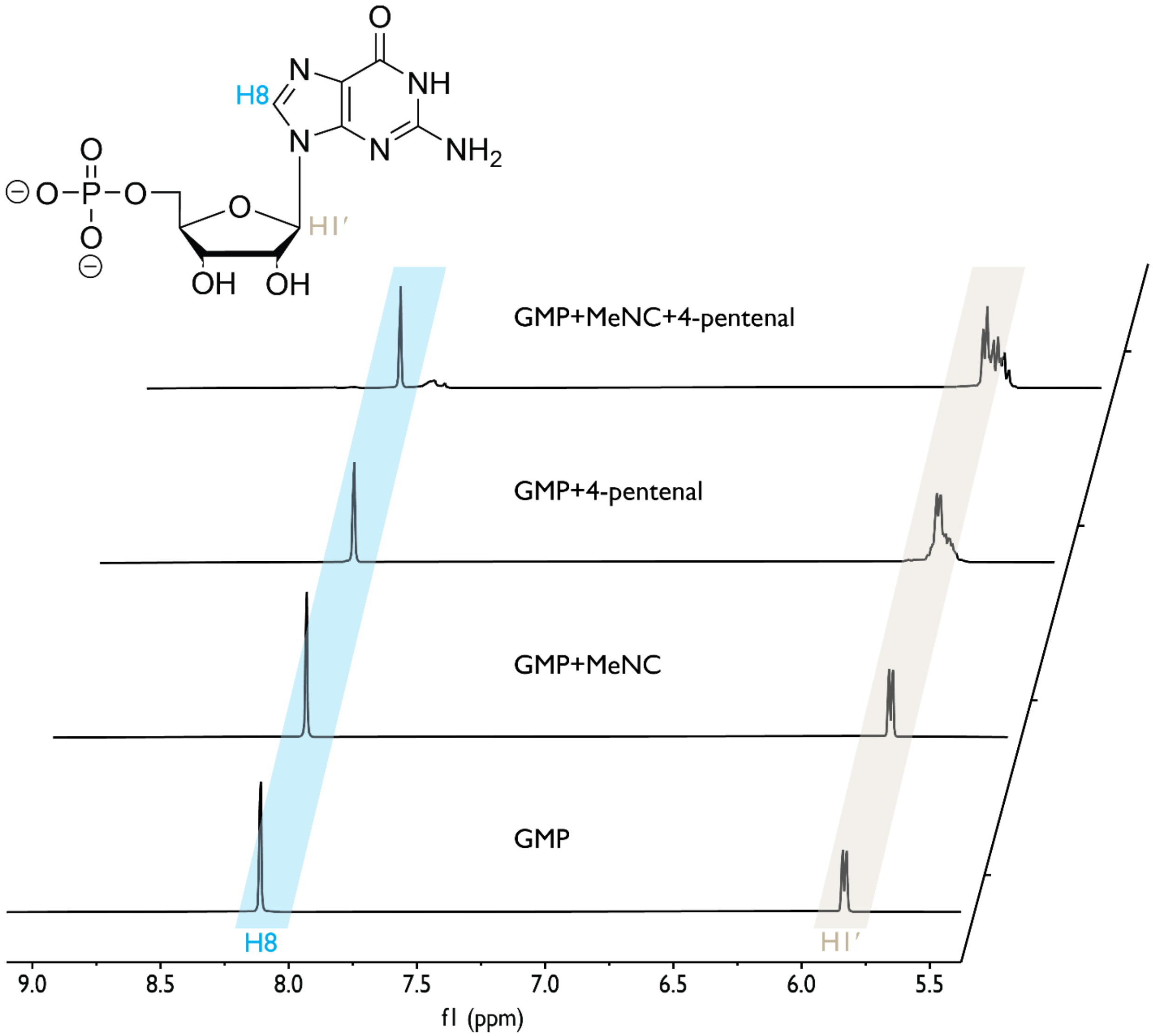


**Figure S8**. Guanosine 5′-monophosphate (GMP) $^1$H NMR spectra acquired in 200 mM HEPES, pH 8.0, in 9:1 (v/v) $H_2O$:$D_2O$ after 24 h. (1) 25 mM GMP; (2) 25 mM GMP, 100 mM MeNC; (3) 25 mM GMP, 100 mM 4-pentenal; (4) 25 mM GMP, 100 mM MeNC, 100 mM 4-pentenal.

**Fig.S9**

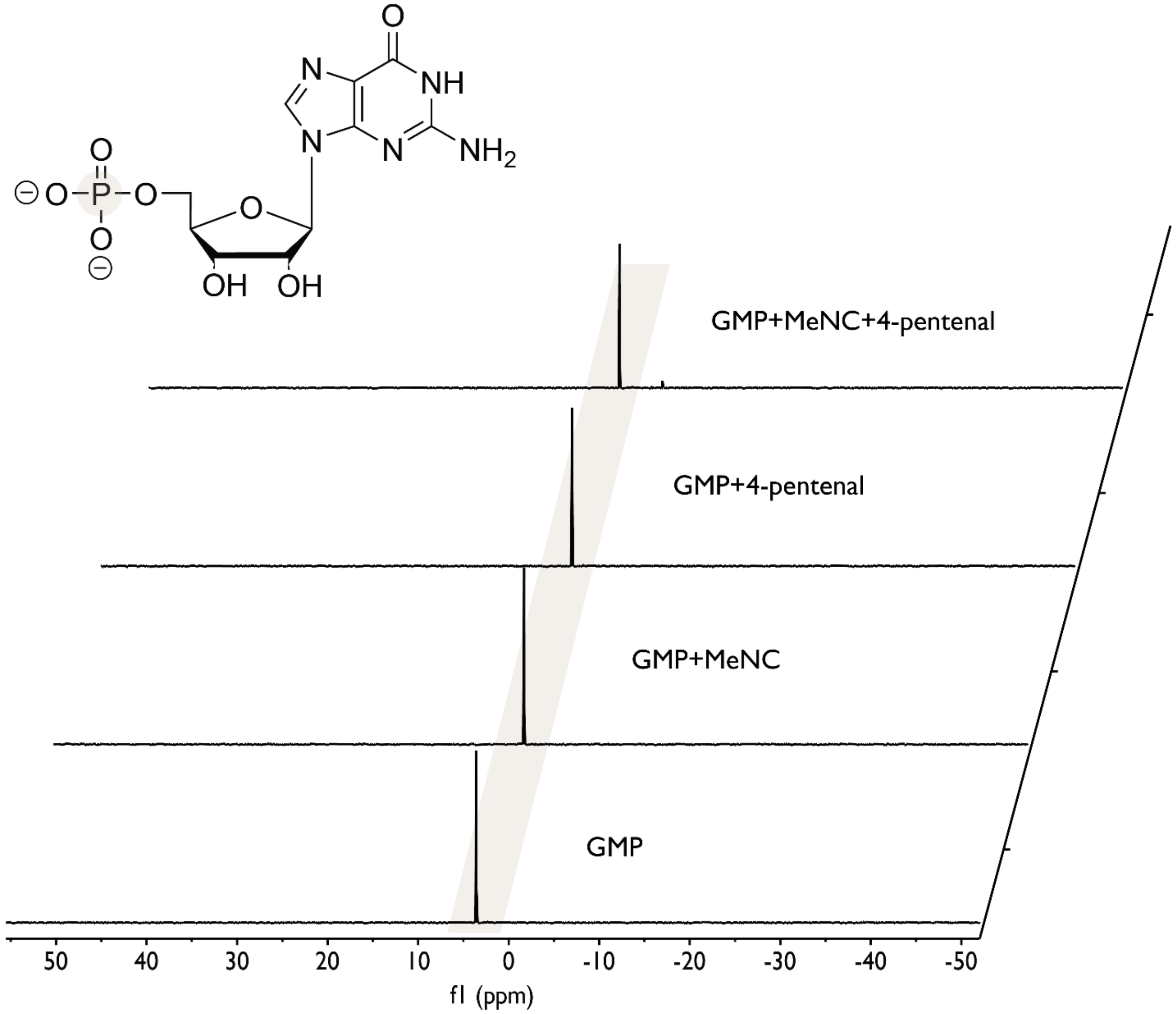


**Figure S9**. Guanosine 5′-monophosphate (GMP) $^{31}P$ NMR spectra acquired in 200 mM HEPES, pH 8.0, in 9:1 (v/v) $H_2O:D_2O$ after 24 h. (1) 25 mM GMP; (2) 25 mM GMP, 100 mM MeNC; (3) 25 mM GMP, 100 mM 4-pentenal; (4) 25 mM GMP, 100 mM MeNC, 100 mM 4-pentenal.

**Fig.S10**

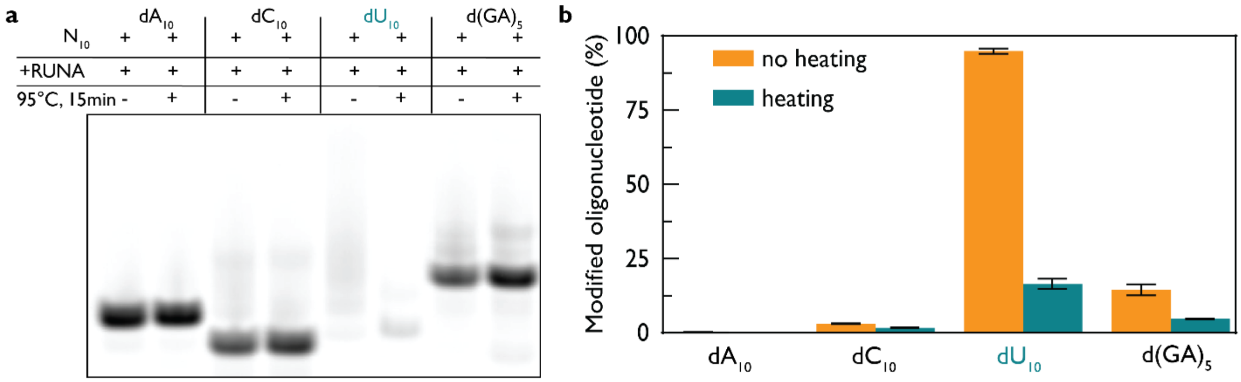


**Figure S10**. 20 % (v/v) denaturing PAGE of 10-mer oligodeoxynucleotides (1 µM $dA_{10}$, $dC_{10}$, $dU_{10}$, and $d(GA)_5$) incubated with 200 mM HEPES, pH 8.0, 200 mM norbornene aldehyde, and 200 mM MeNC. **(a)** Gel indicating the same conditions with and without heating at 95 °C for 15 minutes. **(b)** Quantification of band intensities with error bars representing standard error.

**Fig.S11**

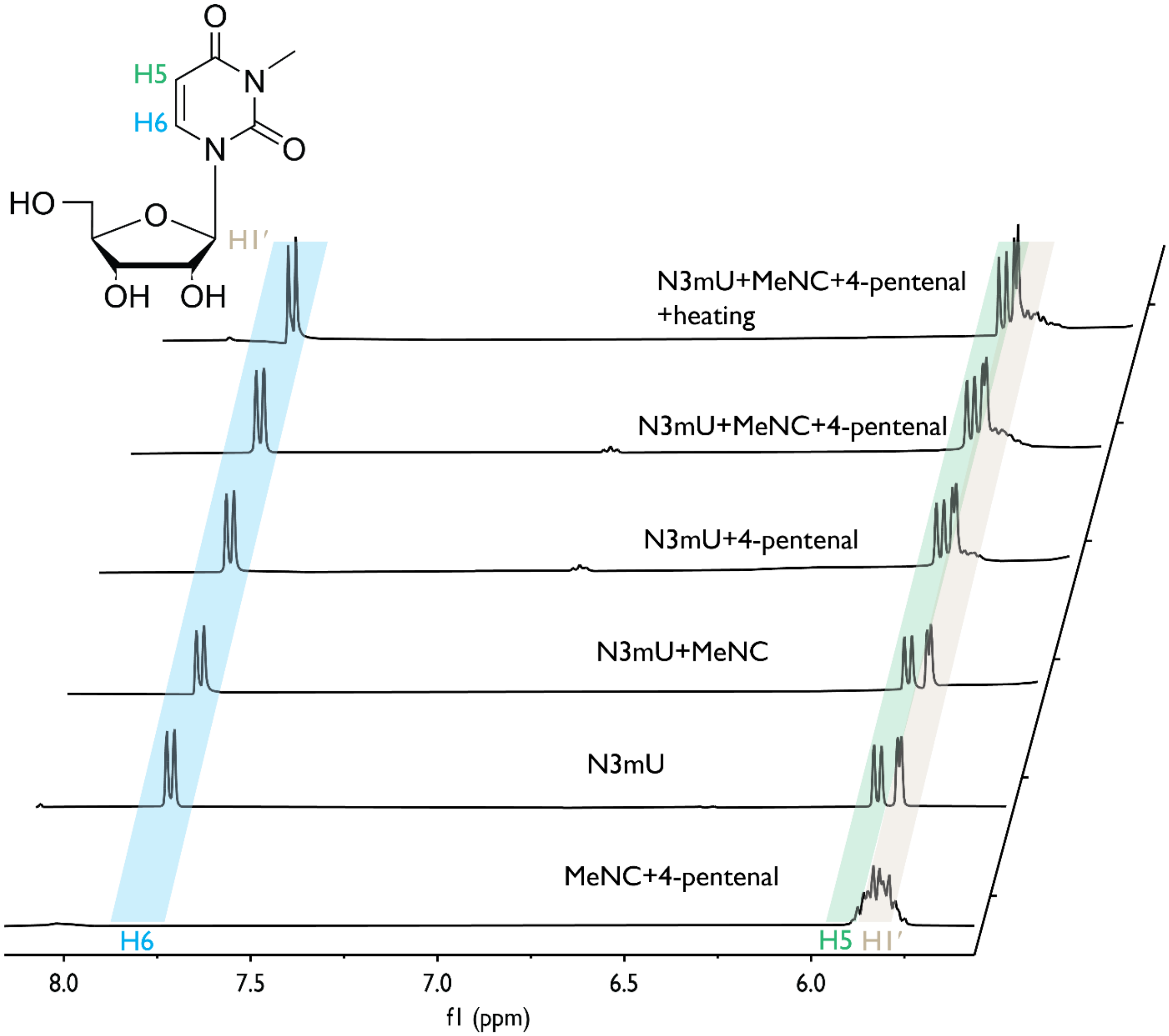


**Figure S11**. N3-methyluridine (N3mU) $^1$H NMR spectra acquired in 200 mM HEPES, pH 8.0, in 9:1 (v/v) $H_2O$:$D_2O$ after 24 h. (1) 100 mM MeNC, 100 mM 4-pentenal; (2) 25 mM N3mU; (3) 25 mM N3mU, 100 mM MeNC; (4) 25 mM N3mU, 100 mM 4-pentenal; (5) 25 mM N3mU, 100 mM MeNC, 100 mM 4-pentenal; (6) sample (5) heated for 15 min at 95 °C.

**Fig.S12**

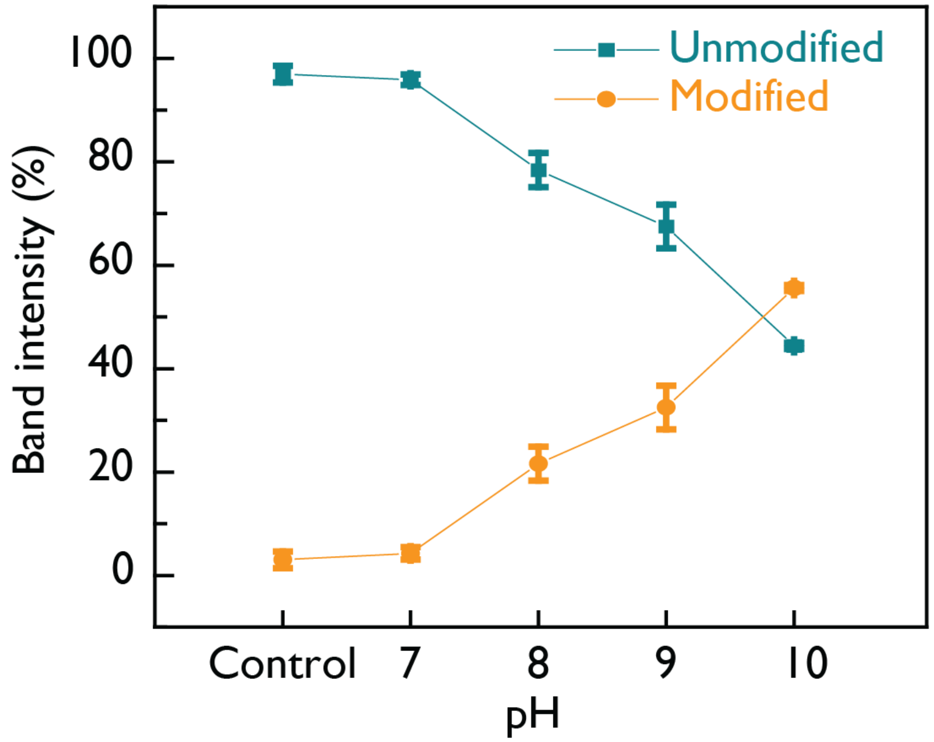


**Figure S12.** 3′ Cy3-labeled 12-mer RNA (CAGCUCUAGA-Cy3) was incubated for 24 h with 200 mM 4-pentenal, 200 mM MeNC, and 200 mM HEPES at pH 7, 8, 9, or 10. Samples were analyzed by denaturing PAGE.

Fig.S13

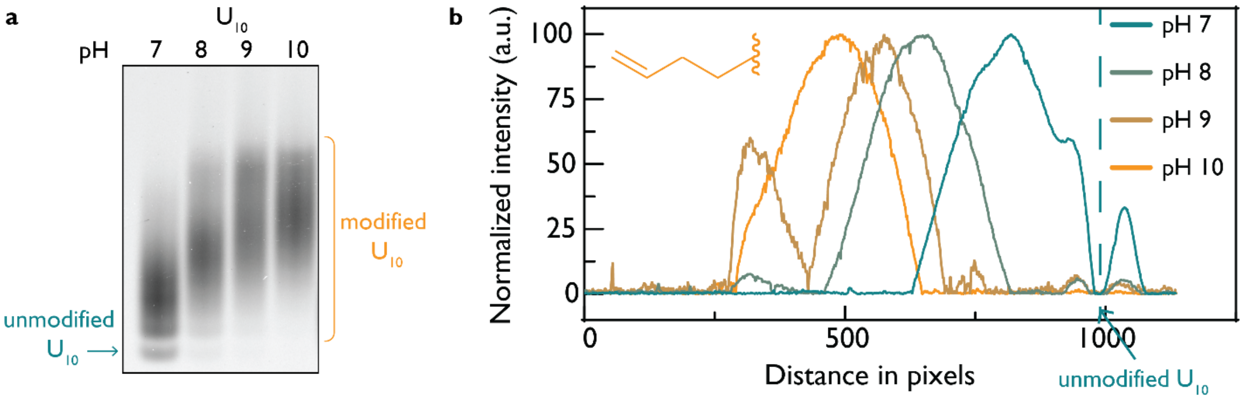


**Figure S13.** 3′ Cy3-labeled $U_{10}$ was incubated for 6 h with 200 mM 4-pentenal, 200 mM MeNC, and 200 mM HEPES at pH 7, 8, 9, or 10, respectively, and then analyzed by 20 % (v/v) PAGE as shown in (**a**) with intensity plots in (**b**). The dashed line separates unmodified (right) from modified (left) $U_{10}$.

**Fig.S14**

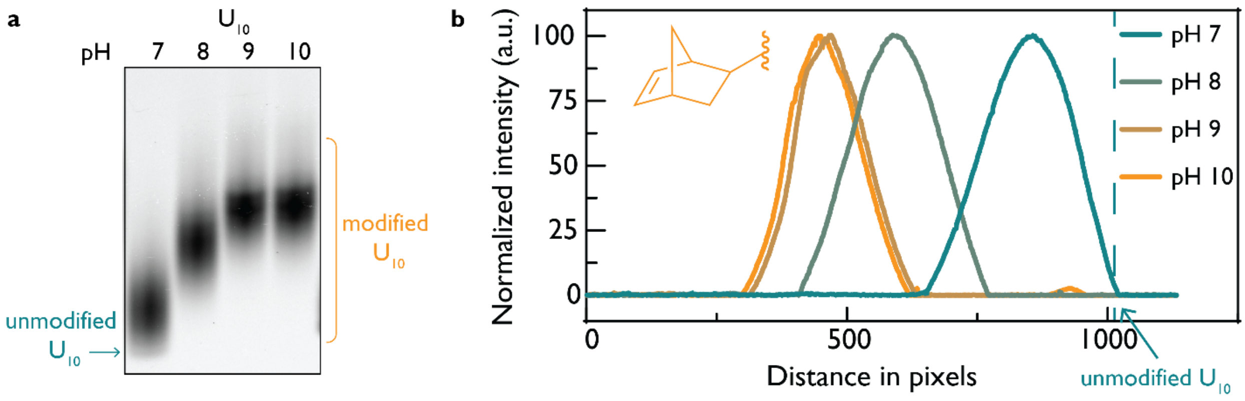


**Figure S14.** 3′ Cy3-labeled $U_{10}$ was incubated for 6 h with 200 mM norbornene aldehyde, 200 mM MeNC, and 200 mM HEPES at pH 7, 8, 9, or 10, respectively, and then analyzed by 20 % (v/v) PAGE as shown in (**a**) with intensity plots in (**b**). The dashed line separates unmodified (right) from modified (left) $U_{10}$.

**Fig.S15**

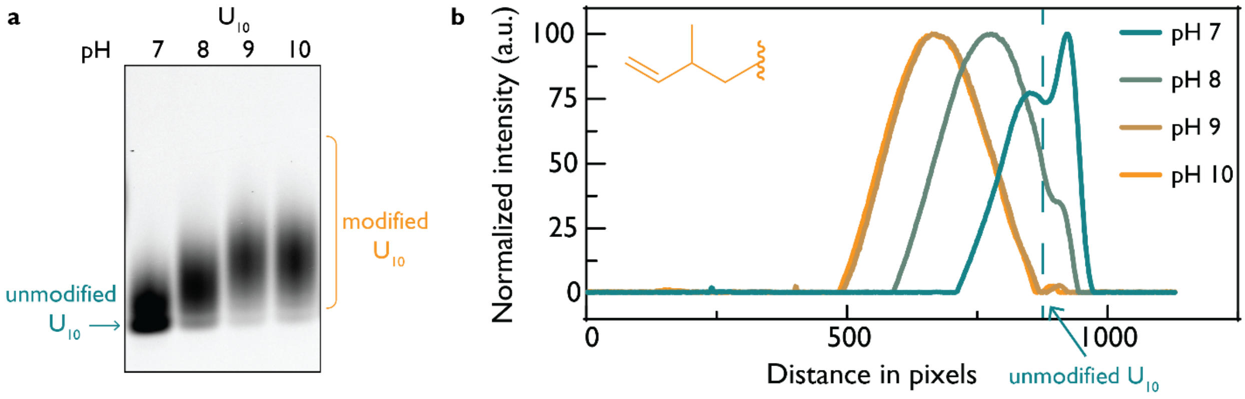


**Figure S15.** 3′ Cy3-labeled $U_{10}$ was incubated for 6 h with 200 mM 2-methyl-4-pentenal, 200 mM MeNC, and 200 mM HEPES at pH 7, 8, 9, or 10, respectively, and then analyzed by 20 % (v/v) PAGE as shown in (**a**) with intensity plots in (**b**). The dashed line separates unmodified (right) from modified (left) $U_{10}$.

**Fig.S16**

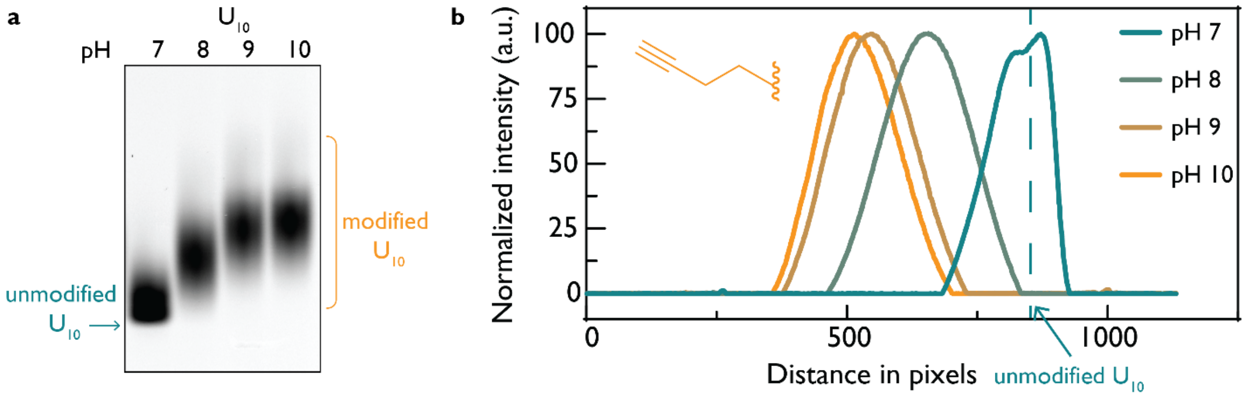


**Figure S16.** 3′ Cy3-labeled $U_{10}$ was incubated for 6 h with 200 mM 4-pentynal, 200 mM MeNC, and 200 mM HEPES at pH 7, 8, 9, or 10, respectively, and then analyzed by 20 % (v/v) PAGE as shown in (**a**) with intensity plots in (**b**). The dashed line separates unmodified (right) from modified (left) $U_{10}$.

**Fig.S17**

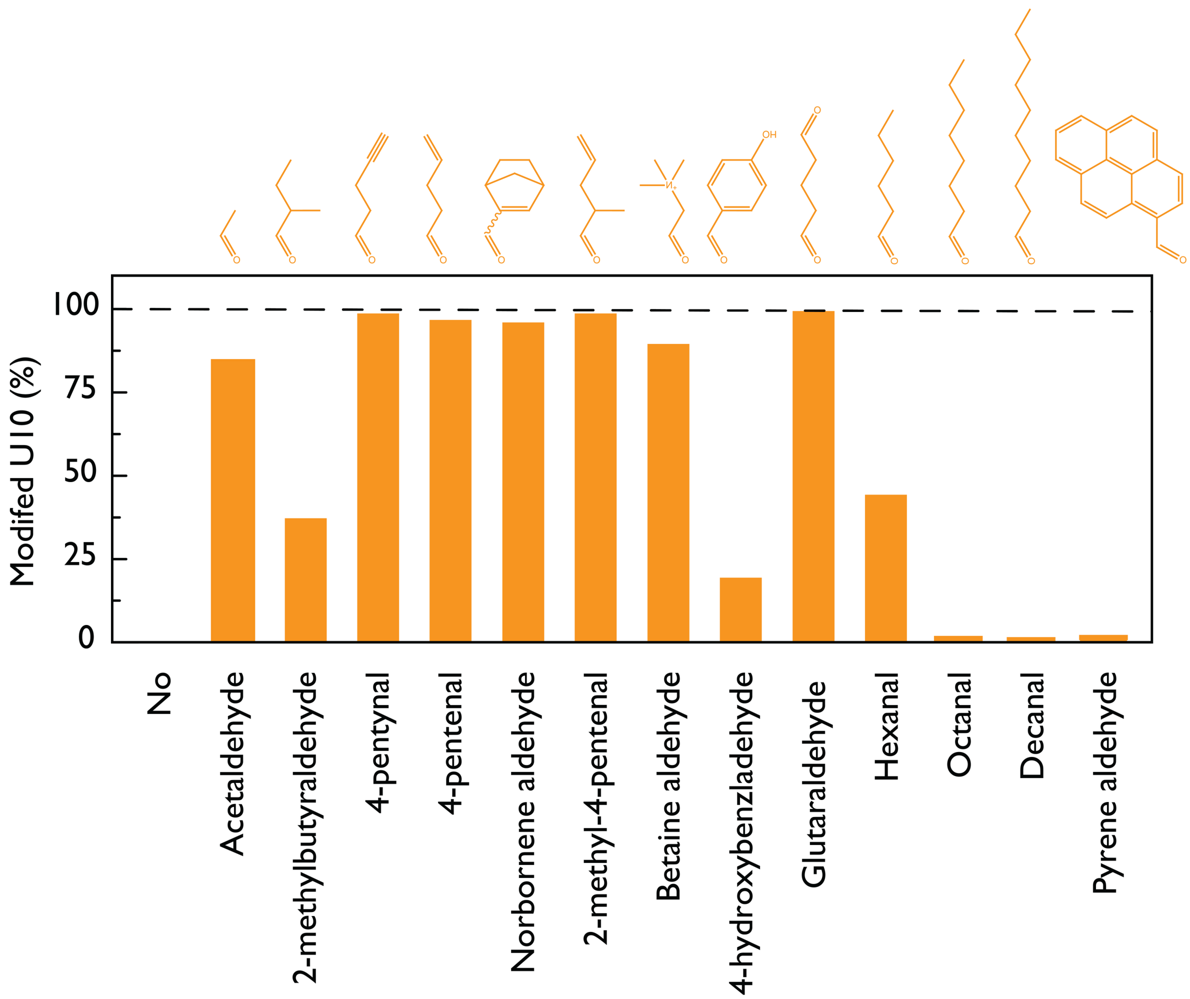


**Figure S17.** 3′ Cy3-labeled $U_{10}$ reactivity with various aldehydes analyzed by 20 % (v/v) PAGE; the percentage of modified $U_{10}$ is reported. Reactions contained 1 μM $U_{10}$, 200 mM aldehyde, 200 mM MeNC, and 200 mM HEPES, pH 8.0, and were incubated for 24 h at 18 °C.

**Fig.S18**

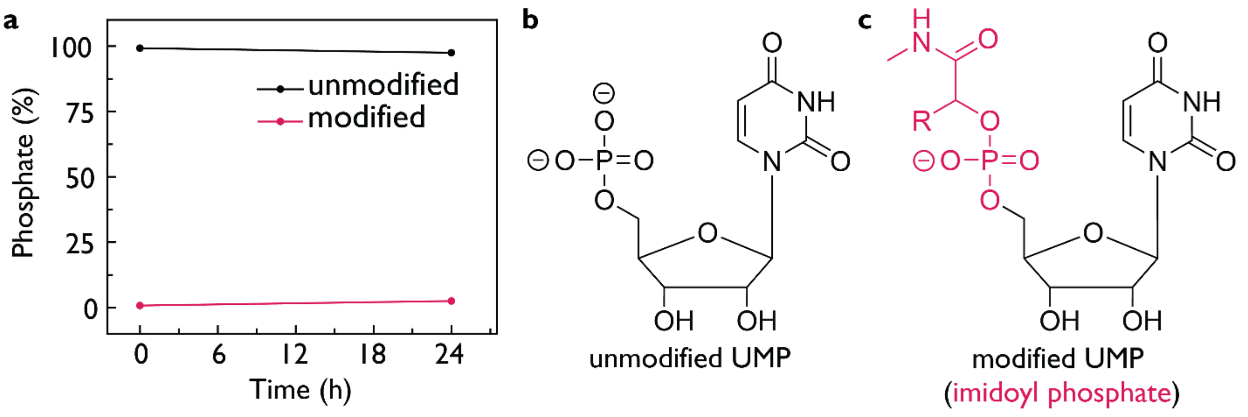


**Figure S18.** $^{31}$P NMR of UMP (25 mM) over time in the presence of 200 mM 4-pentenal, 200 mM MeNC, and 200 mM HEPES, pH 8.0. **(a)** $^{31}$P NMR showing native phosphate and a weak signal consisstent with a imidoyl phosphate adduct even after 24 h. **(b)** Structure of UMP; **(c)** proposed UMP-imidoyl phosphate structure.

**Fig.S19**

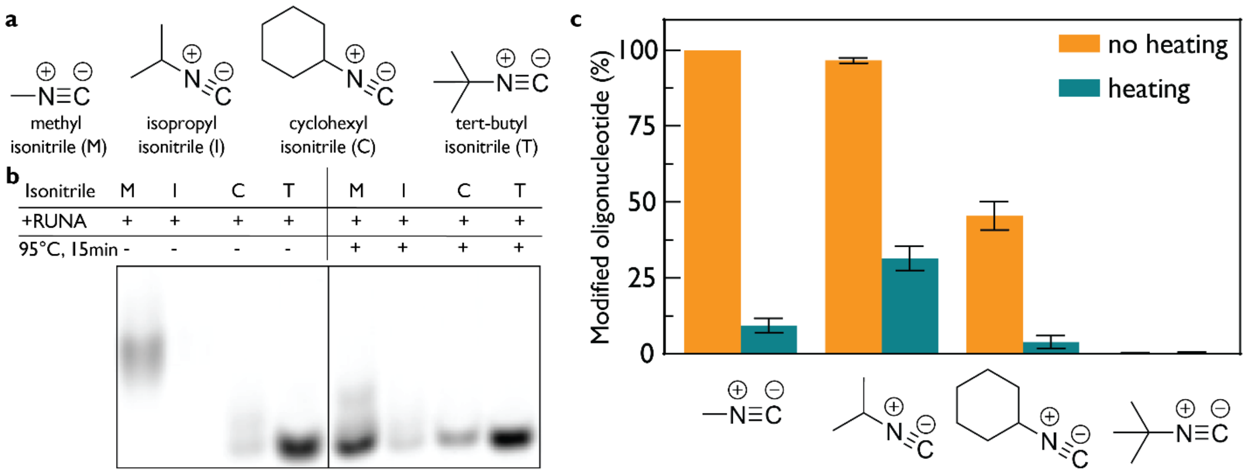


**Figure S19.** 3′ Cy3-labeled 12-mer RNA (1 μM) was incubated for 24 h with 200 mM norbornene aldehyde, 200 mM of the indicated isonitrile, and 200 mM HEPES, pH 8.0. **(a)** Four isonitriles tested. **(b)** 20 % (v/v) PAGE before and after heat-induced reversal (95 °C, 15 min): Lane 1, MeNC; Lane 2, isopropyl isonitrile; Lane 3, cyclohexyl isonitrile; Lane 4, *tert*-butyl isonitrile. **(c)** Quantification of band intensities before and after heat-induced reversal with error bars representing standard error.

**Fig.S20**

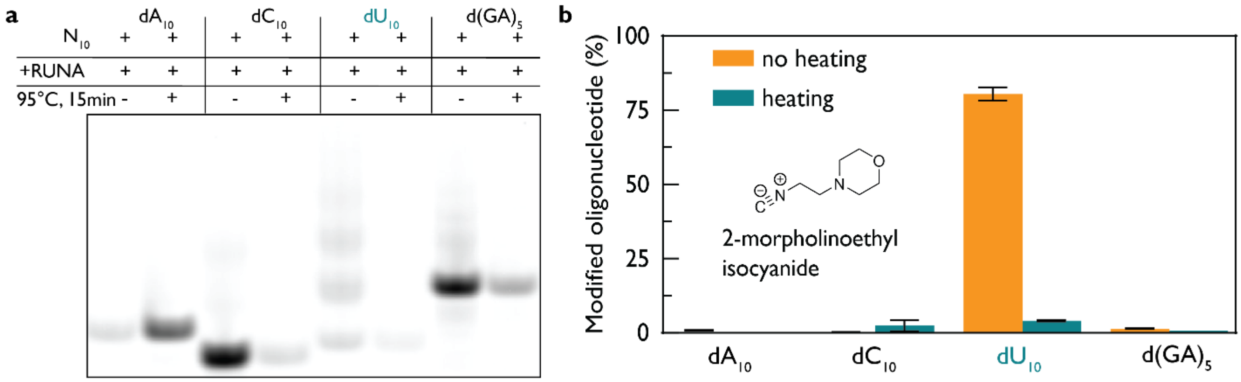


**Figure S20.** 3′ Cy3-labeled $dN_{10}$ was incubated for 6 h with 200 mM norbornene aldehyde, 200 mM 2-morpholinoethyl isonitrile, and 200 mM HEPES, pH 8.0, then analyzed by **(a)** 20 % (v/v) PAGE before and after heat-induced reversal (95 °C, 15 min). **(b)** Quantification of band intensities before and after heat-induced reversal with error bars representing standard error.

**Fig.S21**

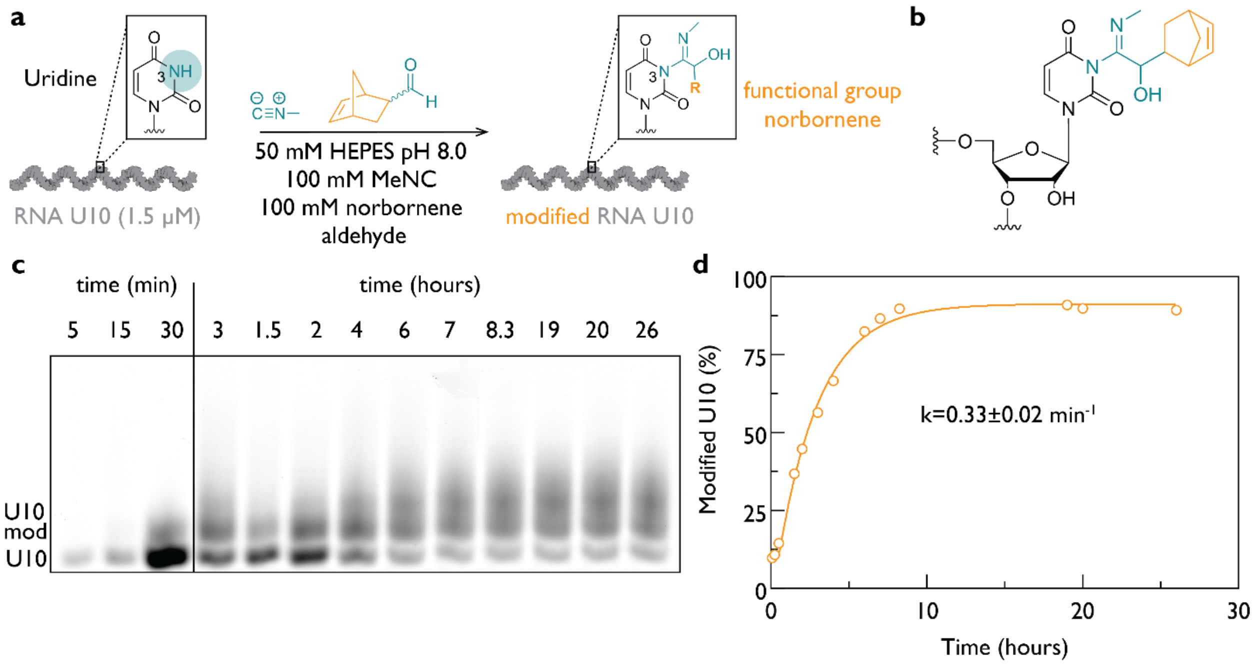


**Figure S21.** 3′ Cy3-labeled $U_{10}$ RNA (1.5 μM) was incubated with 100 mM MeNC and 100 mM norbornene aldehyde in 50 mM HEPES, pH 8.0, over time. **(a)** Reaction scheme; **(b)** structure of the norbornene adduct of N3 of uridine in modified $U_{10}$; **(c)** denaturing gel showing time-dependent modification; **(d)** kinetic fit giving k = 0.33 ± 0.02 $min^{-1}$ (GraphPad Prism 10).

**Fig.S22**

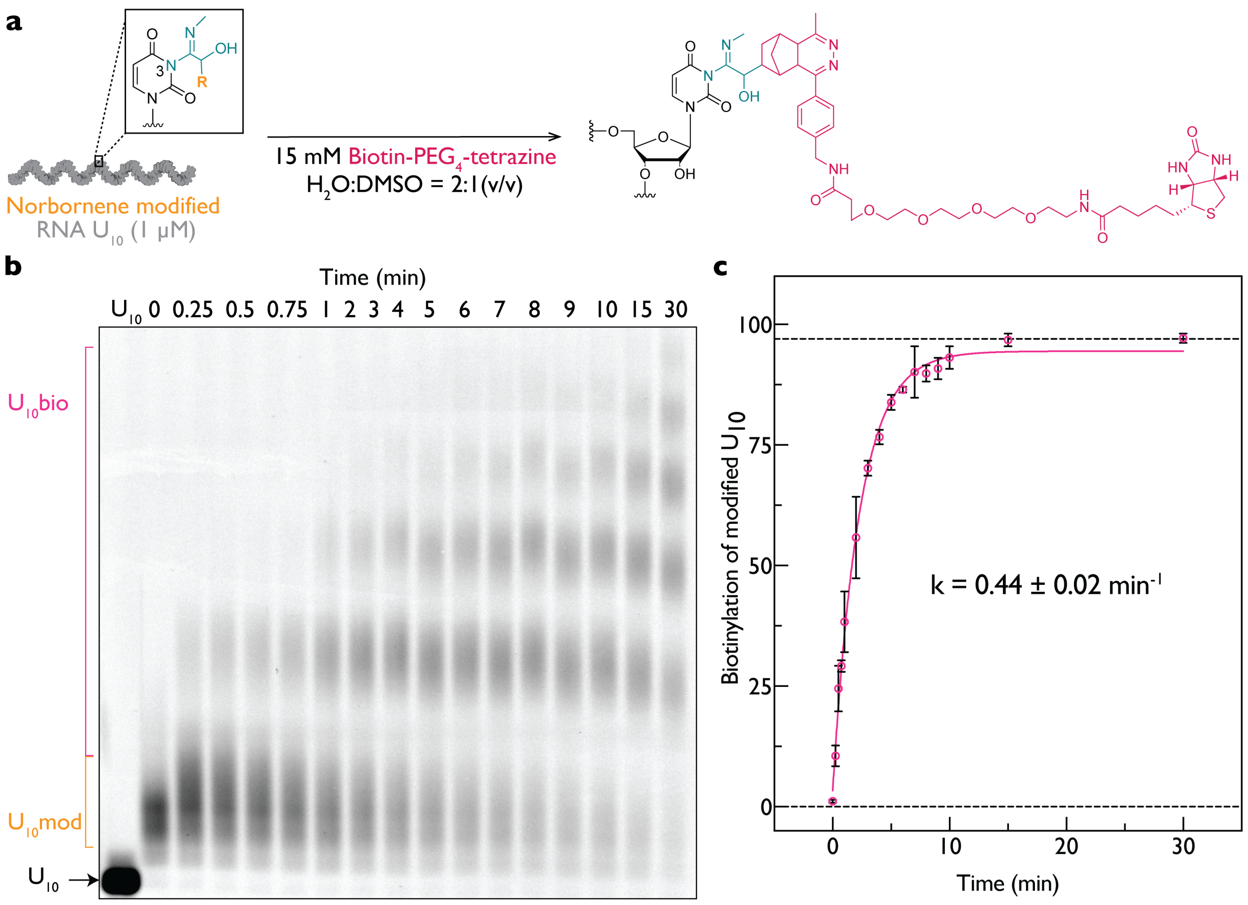


**Figure S22. Kinetics of labeling of norbornene-modified 3′-Cy3-labeled U₁₀ with biotin-PEG₄-tetrazine.** (**a**) Reaction of norbornene-modified, 3′-Cy3-labeled $U_{10}$ RNA with biotin-$PEG_4$-tetrazine (Lumiprobe). (**b**) A 20% (v/v) PAGE analysis shows time-dependent formation of the biotin-labeled $U_{10}$ product during incubation with 15 mM biotin-$PEG_4$-tetrazine (Lumiprobe) in $H_2O$:DMSO (2:1, v/v). Reactions were quenched by adding 0.5 µL of biotin-$PEG_4$-labeled $U_{10}$ RNA directly to a premixed solution containing 8.5 µL of 98% (v/v) formamide, 5 mM EDTA, and 0.5 µL of 500 mM (E)-cyclooct-4-enol (TCO-OH). The mixture was vortexed, and 1 µL was loaded onto the gel. (**c**) An apparent rate constant of 0.44 ± 0.02 $min^{-1}$ was obtained by fitting the data in GraphPad Prism 10 to a single exponential model. Data represent five replicates.

**Fig.S23**

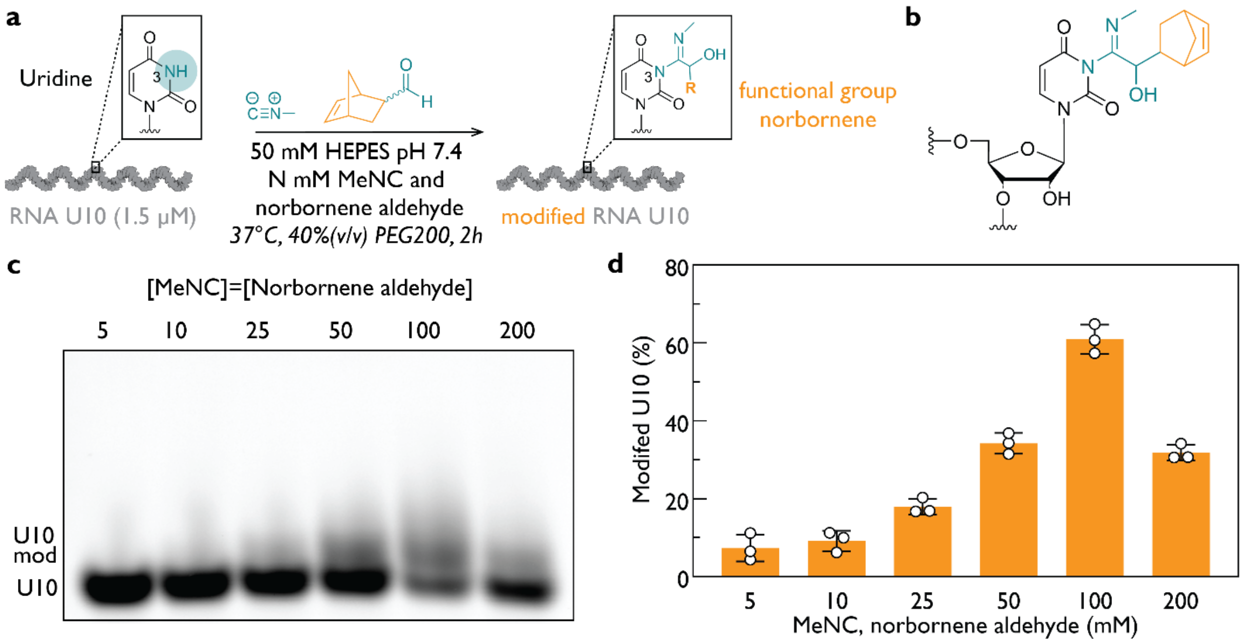


**Figure S23.** 3′ Cy3-labeled $U_{10}$ RNA (1.5 µM) was incubated in 50 mM HEPES, pH 7.4, with 40 % (v/v) PEG-200 at 37 °C in the presence of MeNC and norbornene aldehyde at the indicated concentrations for 2 h. **(a)** Reaction scheme and the conditions used; **(b)** structure of adduct; **(c)** 20 % (v/v) PAGE showing modified $U_{10}$ with varying concentrations of MeNC and norbornene aldehyde; **(d)** Quantification of band intensities from (c) with error bars representing standard error.

Fig.S24

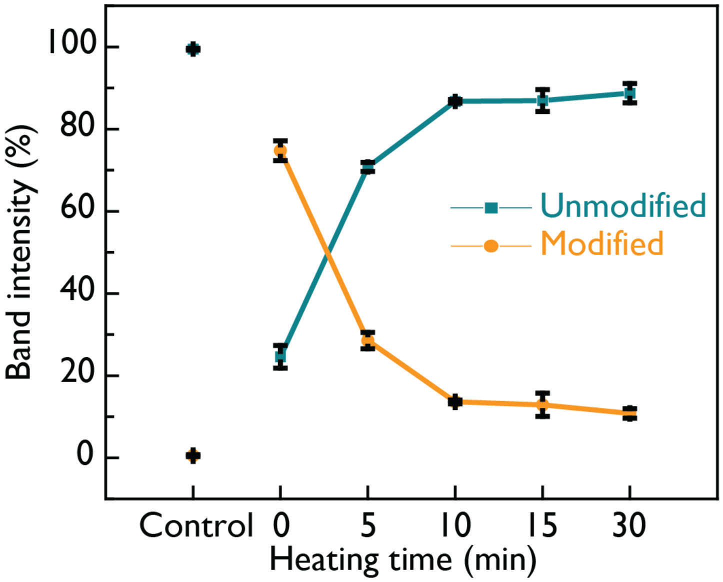


**Figure S24.** 3′ Cy3-labeled 12-mer RNA was incubated for 12 h with 200 mM 4-pentenal, 200 mM MeNC, and 200 mM HEPES, pH 8.0, then heated for 0–30 min at 95 °C and analyzed by 20 % (v/v) PAGE. Quantification of band intensities of unmodified and modified 12-mer RNA at 0 min, 5 min, 10 min, 15 min, and 30 min, respectively, is shown with error bars representing standard error.

**Fig.S25**

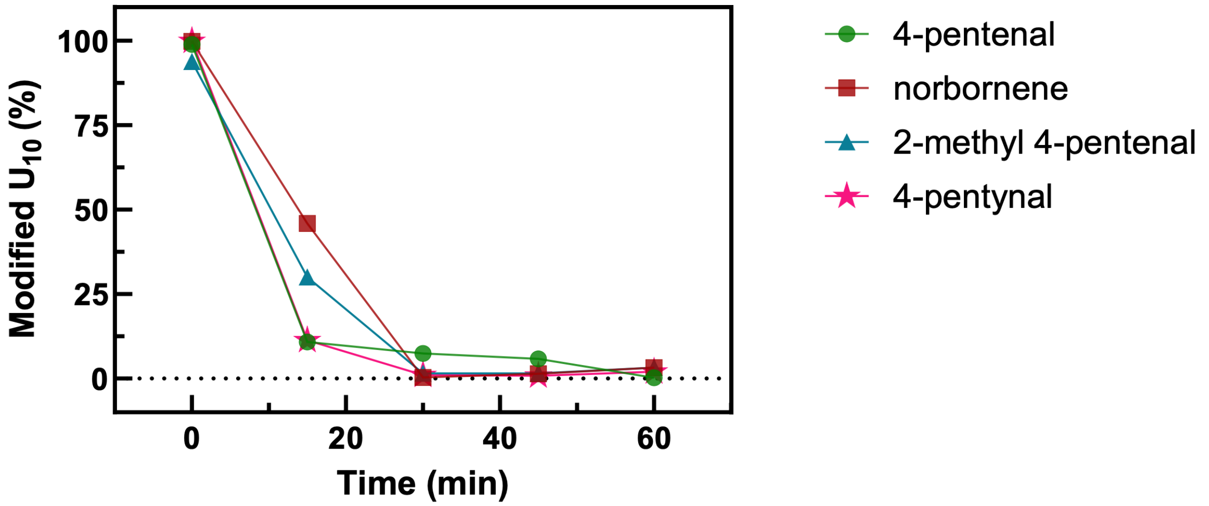


**Figure S25.** Reversible removal of 3′ Cy3-labeled $U_{10}$ RNA modifications formed with 4-pentenal, norbornene, 2-methyl-4-pentenal, or 4-pentynal (200 mM aldehyde, 200 mM MeNC, and 200 mM HEPES, pH 8.0, after 24 h). Modified samples were heated at 95 °C for indicated times and analyzed by 20 % (v/v) PAGE.

**Fig.S26**

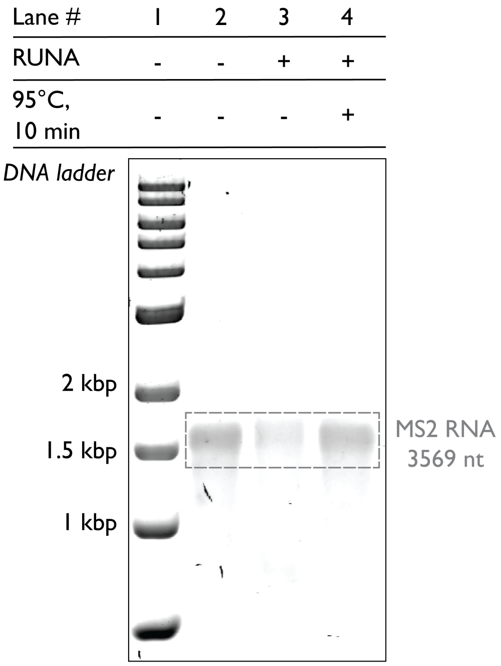


**Figure S26. Native MS2 RNA genome (3,569 nt, Roche) RUNA labeling and reversal**. 1% (w/v) agarose gel in 1× TBE, 4 °C. Lanes: (1) dsDNA ladder (New England Biolabs): 1, 1.5, and 2 kbp. (2) Unmodified MS2 RNA. (3) MS2 RNA labeled *via* RUNA using MeNC and norbornene aldehyde (100 mM each, 2 h, 200 mM HEPES, pH 8.0). (4) Sample from lane 3 heated at 95 °C for 10 min. The gel was post-stained with 1× SYBR Gold (Invitrogen) in 1× TBE for 10 minutes.

**Fig.S27**

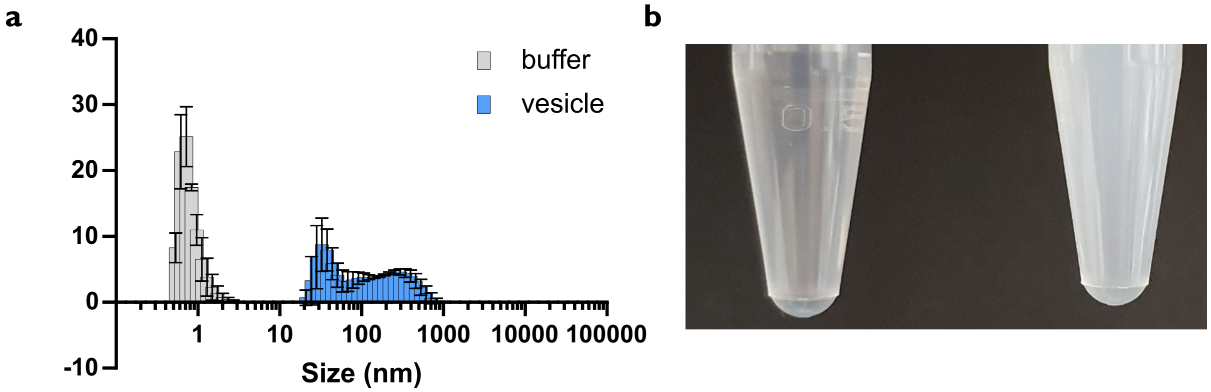


**Figure S27. (a)** DLS particle-size distribution of buffer and vesicle fractions after SEC showing vesicle-associated peaks and **(b)** visible turbidity only in the vesicle fraction.

**Fig.S28**

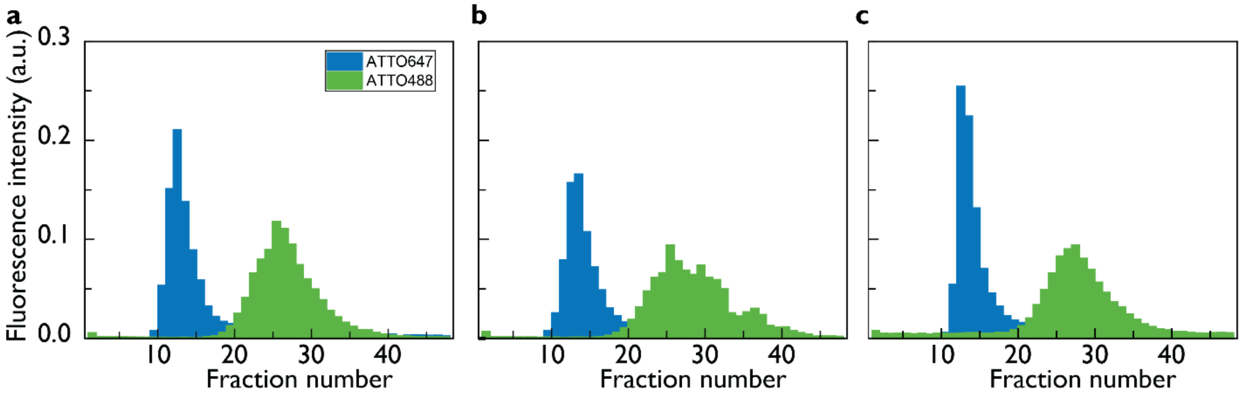


**Figure S28.** SEC of extravesicular and intravesicular RNA of identical 12-mer RNA sequence labeled with ATTO488 and ATTO647 (**Table S1**) after incubation with 200 mM MeNC, 200 mM aldehyde in 200 mM HEPES at pH 8.0. Chromatograms are shown for the following aldehydes used **(a)** acetaldehyde, **(b)** norbornene aldehyde, and **(c)** betaine aldehyde.

**Fig.S29**

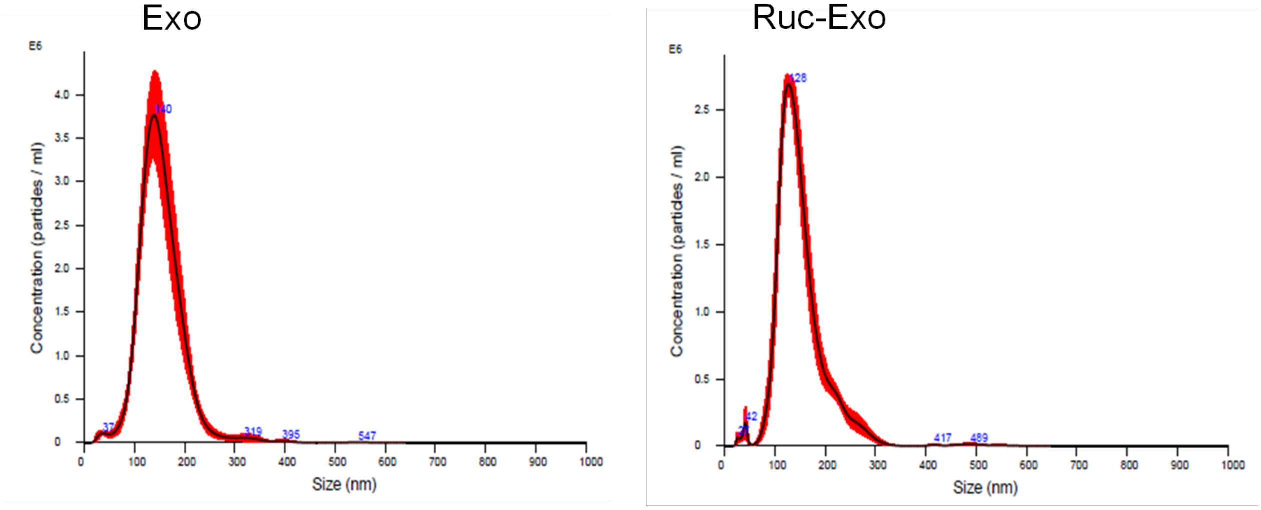


**Figure S29.** NTA of exosomes from control and rucaparib-treated cells showing that exosome size distribution remained unchanged following rucaparib treatment, as shown by the NTA analysis (NanoSight). Once the concentration of particles was determined, equal numbers of exosomes from each condition were used for the experiments. The average concentration of vesicles for each condition from three independent measurements is $3.32 \times 10^8$ vesicles/mL (Exo) and $2.14 \times 10^8$ vesicles/mL (Ruc-Exo).

**Fig.S30**

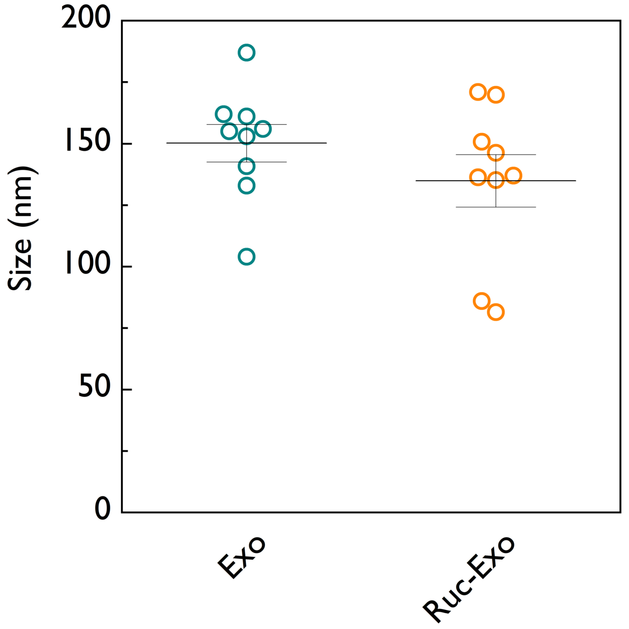


**Figure S30.** DLS of exosomes from control and rucaparib-treated cells showing independently that vesicle sizes remained within the expected range for exosomes (30–200 nm) following rucaparib treatment, as observed with NTA (Figure S29).

**Fig.S31**

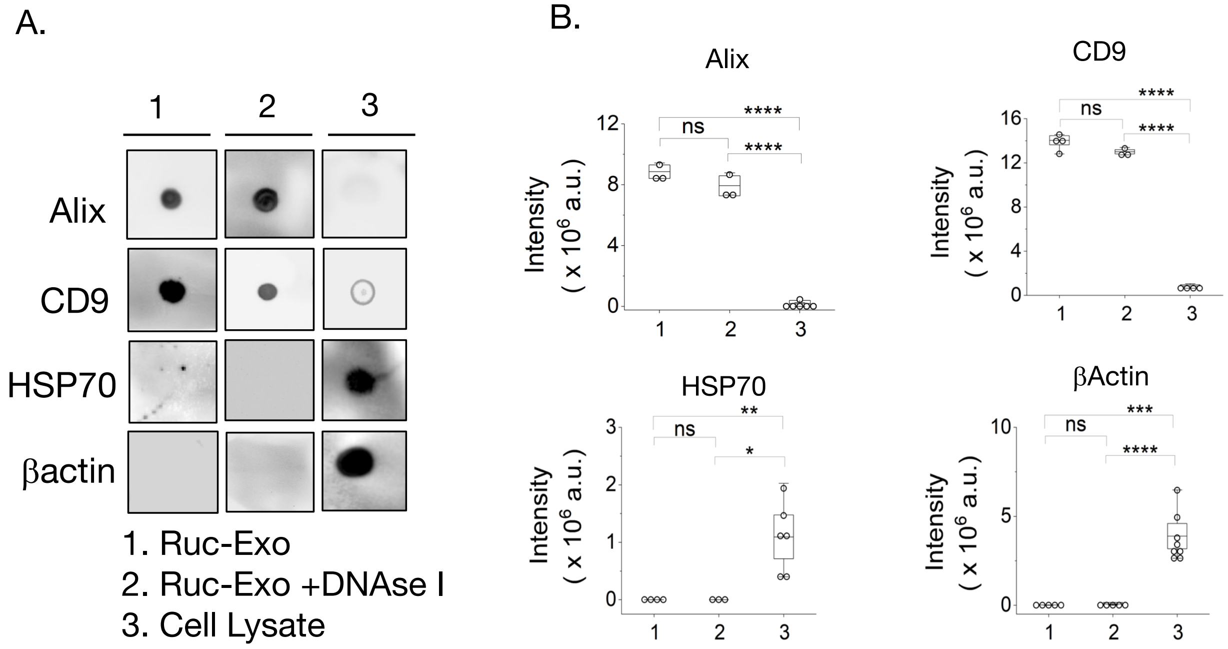


**Figure S31.** DNAse I-treated and untreated exosome fractions exhibit comparable levels of canonical exosome markers CD9 and ALIX, with concomitant reduction of cellular β-actin and HSP70. (A) representative dot blot images for groups 1-3, for exosomal markers (CD9, Alix) and cell specific proteins (HSP70, β-actin); (B) Quantification of the indicated proteins across groups 1-3 from dot blots; plots represent the mean ± SEM, n= 3-8 technical replicates per group from N=3 biological replicates.

**Fig.S32**

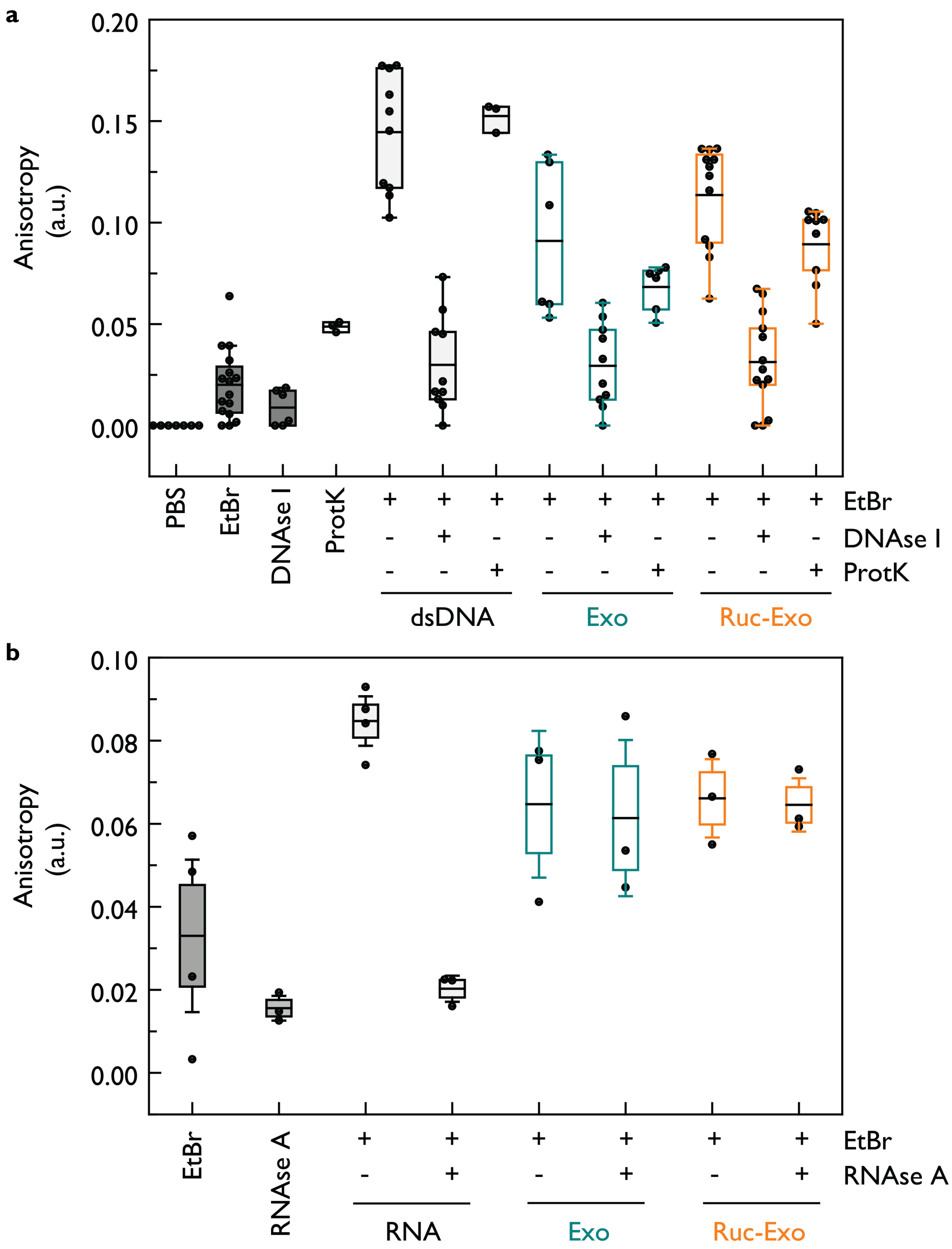


**Figure S32.** Fluorescence polarization (anisotropy) assay using ethidium bromide to detect surface-accessible nucleic acids on exosomes. Equal numbers of exosome pellets from vehicle- and rucaparib-treated samples, along with dsDNA/RNA controls, were subjected to either **(a)** Shrimp DNase treatment (37 °C, 30 min), Proteinase K treatment (37 °C, 30 min); **(b)** RNAase A treatment (37 °C, 30 min) or no treatment. Samples were subsequently stained with 2.5-5 μM ethidium bromide for 30 min at room temperature, and fluorescence anisotropy was measured using a fluorometer. The cumulative results show the relative contributions of surface DNA and proteins to ethidium bromide binding.

Fig.S33

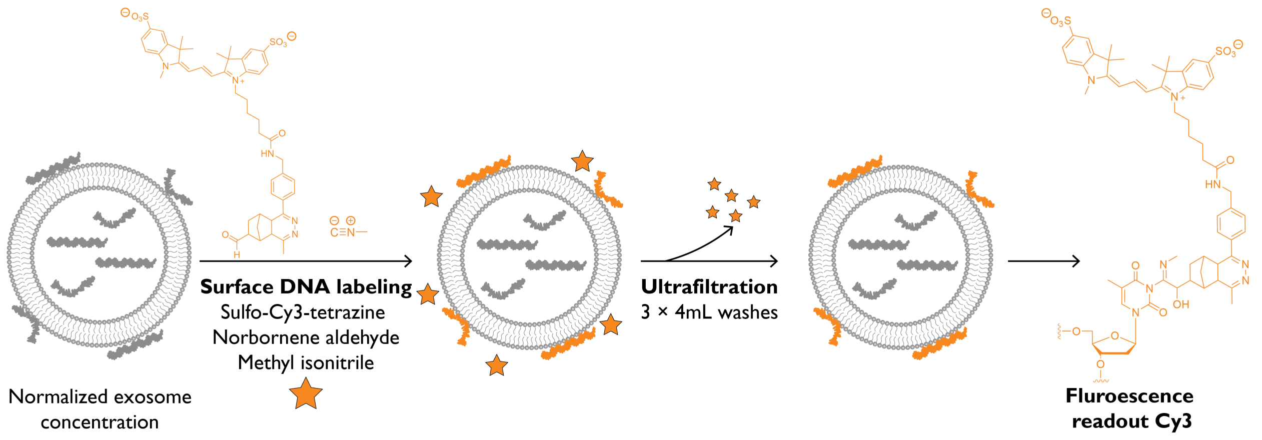


**Figure S33.** Exosomes with the same number of particles, as calculated by NTA (40–60 µL), were surface-labeled by RUNA chemistry using 100 mM norbornene aldehyde and 100 mM MeNC, followed by reaction with 5 mM sulfo-Cy3 tetrazine (Lumiprobe). Purified exosomes were washed (Amicon 4 mL, 100 kDa cutoff) and analyzed by fluorometry (excitation 498 nm; emission 520–650 nm).

**Fig.S34**

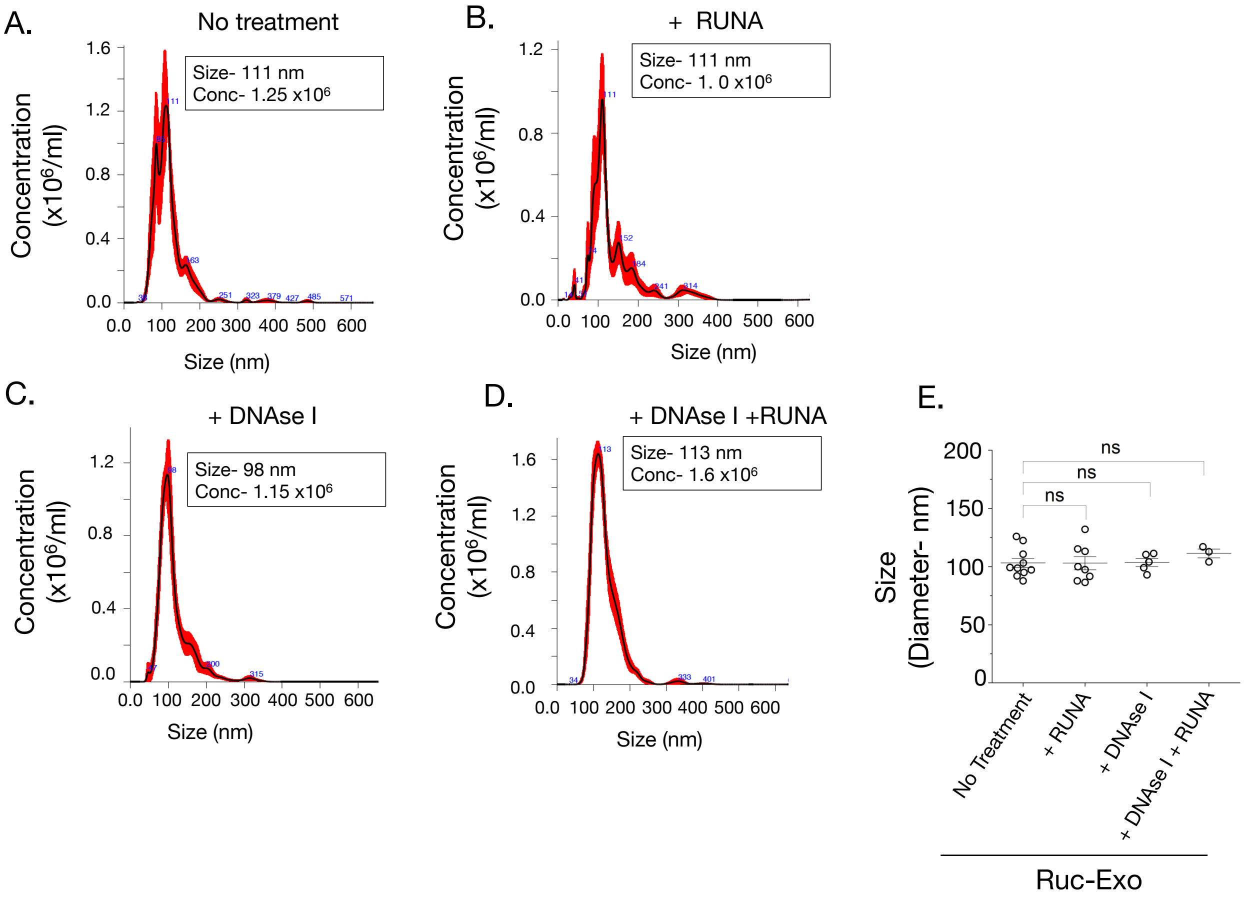


**Figure S34.** Vesicle size is unaffected by *DNase I treatment and RUNA labeling.* (A-D) Representative size distribution and concentration, as given by nanoparticle tracking analysis, of the exosome fraction across untreated and indicated treatment groups. (E) Quantification of exosome sizes in (A-D); data points represent the mean ± SEM, n=3-10 technical replicates obtained from N=2-3 biological replicates.

**Fig.S35**

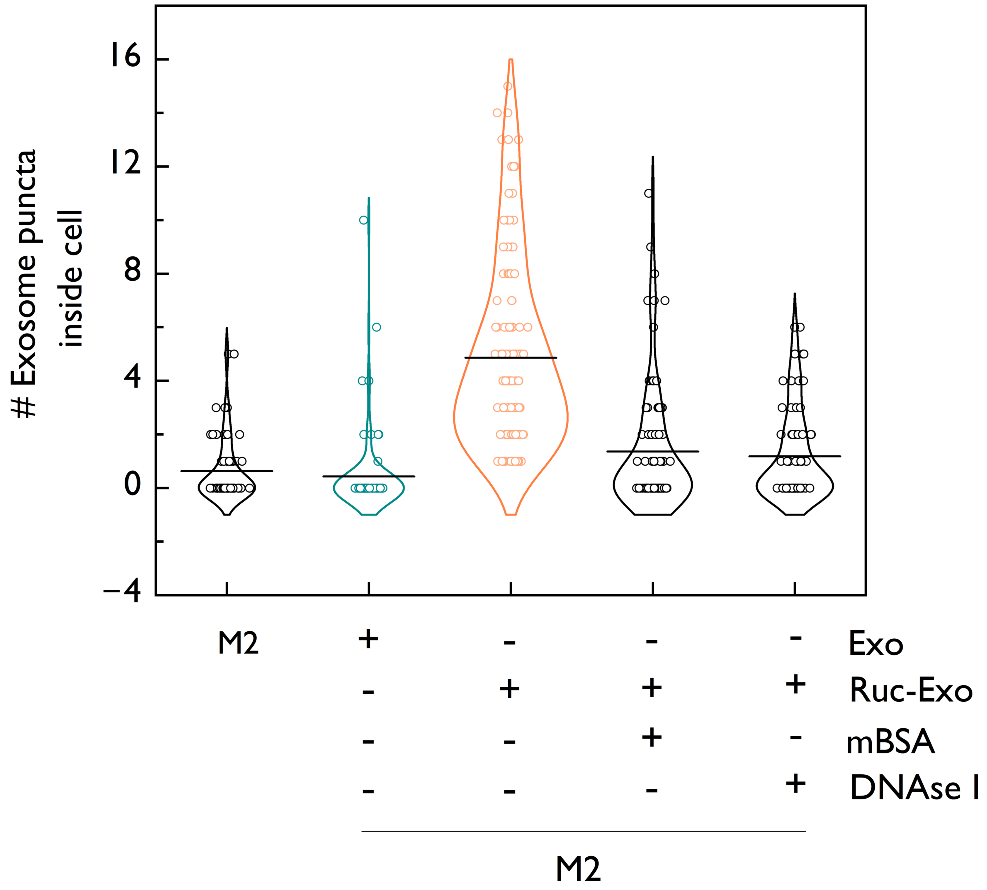


**Figure S35.** Quantification of exosome uptake by M2 macrophages under different treatment conditions. M2 cells were left untreated or treated with exosomes (Exo), rucaparib-treated exosomes (Ruc-Exo), or Ruc-Exo pretreated with maleylated bovine serum albumin (mBSA) and DNase I. The number of exosome particles internalized per cell was calculated to assess the effect of rucaparib treatment and surface-DNA removal on exosome uptake efficiency; n = 70–120 cells.

**Fig.S36**

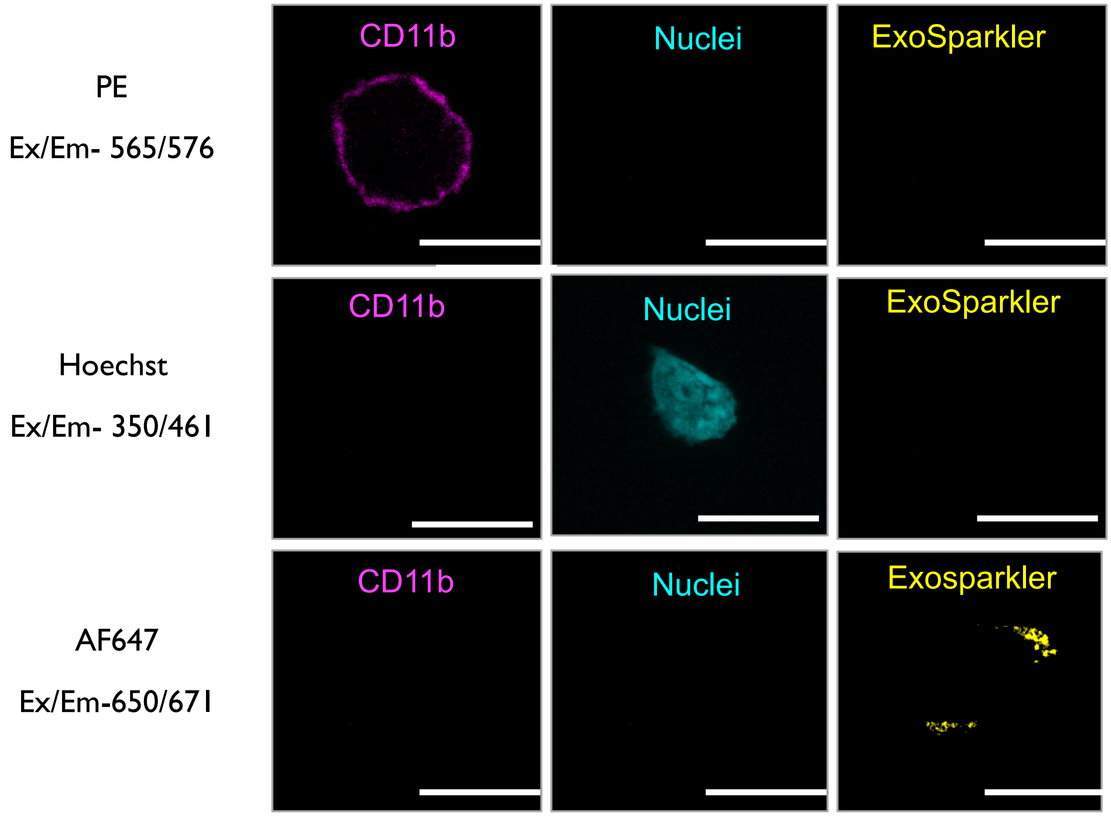


**Figure S36.** Confocal fluorescence imaging showing no spectral overlap between detection channels. Nuclei were stained with Hoechst (405 nm), macrophage surface marker CD11b was detected using PE (561 nm), and exosomes were labeled with ExoSparkler AF647 (640 nm). Distinct fluorescence signals confirm clear spectral separation between channels and reliable detection of exosome uptake in $CD11b^+$ macrophage cells. The scale bar is 10 µm.

**Fig.S37**

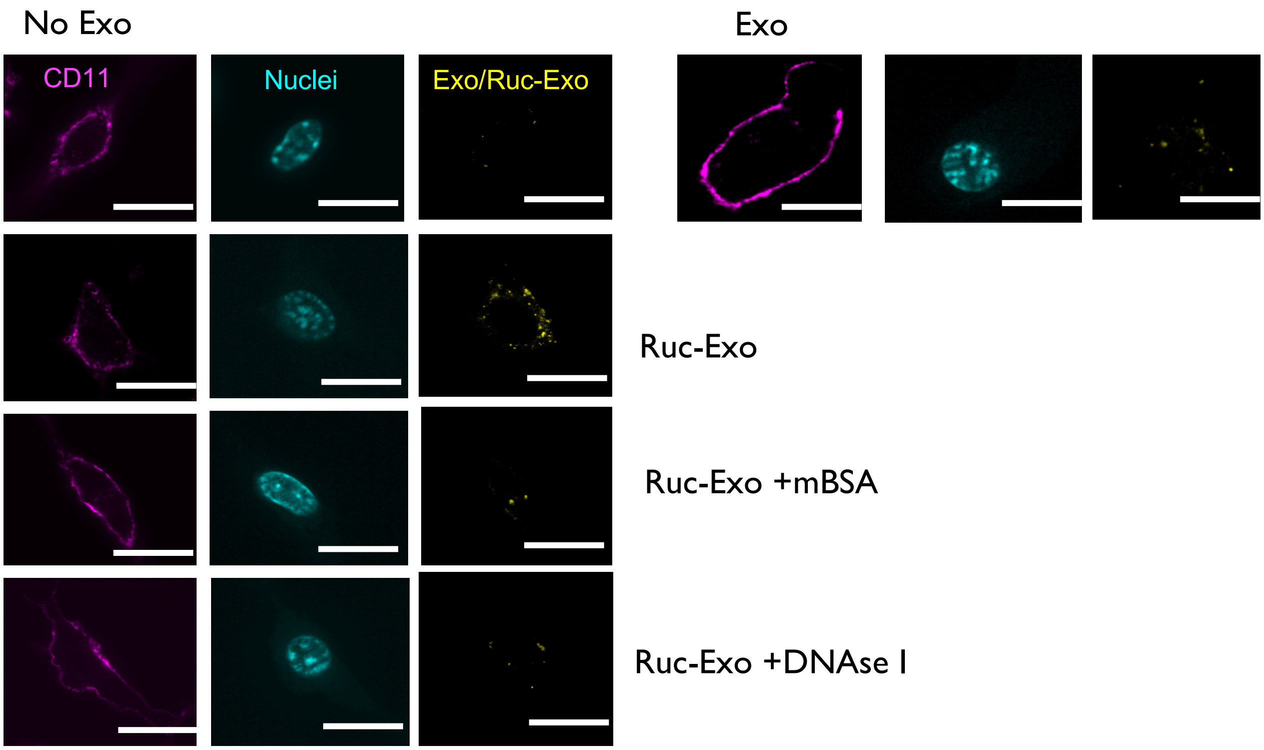


**Figure S37.** Individual fluorescence channels corresponding to the main representative images used for quantification of exosome uptake by M2 macrophages. M2 cells were either untreated or treated with exosomes (Exo), rucaparib-treated exosomes (Ruc-Exo), or Ruc-Exo pretreated with maleylated bovine serum albumin (mBSA) and DNase I. Separate channels display Hoechst-stained nuclei (blue), CD11b$^{+}$ macrophage membrane marker (orange), and ExoSparkler AF647–labeled exosomes (red). The scale bar represents 10 µm.

**Fig.S38**

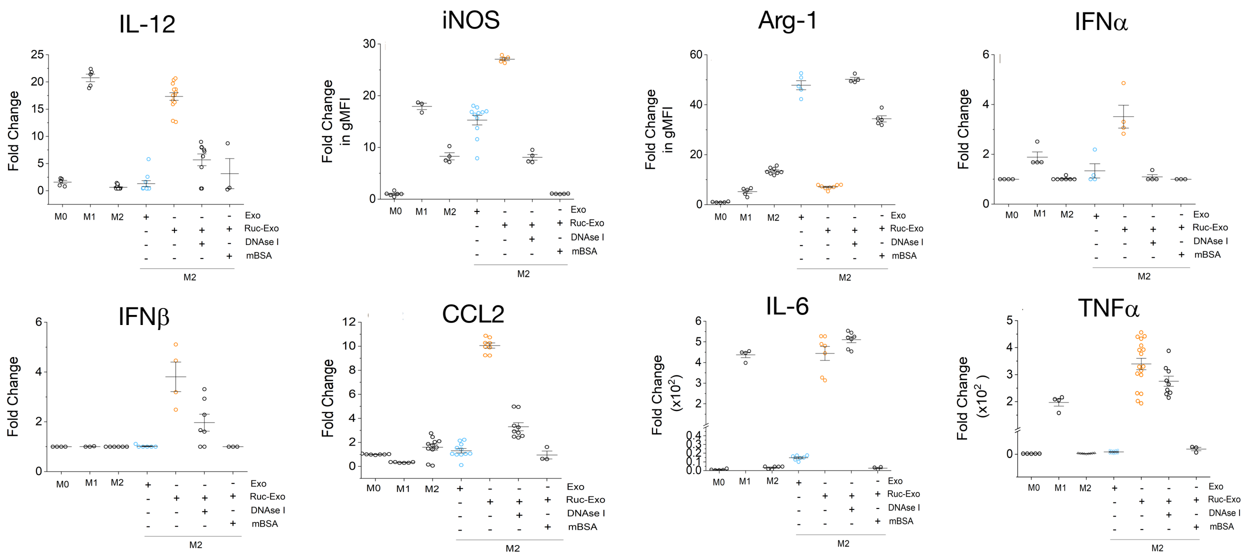


**Figure S38.** Supernatants of M2 macrophages treated with the indicated reagents analyzed by flow cytometry for the indicated cytokines, chemokines and intracellular markers. M2 macrophages were incubated with either unlabeled Ruc-Exo (exosomes from drug-treated cancer cells) or Exo (exosomes from untreated cancer cells). Differential expression of cytokines/chemokines, including IFNβ, IFNα, IL-12, IL-6, CCL2, and TNF-α (concentration in pg/ml), as well as early intracellular markers iNOS and Arg-1 (flow cytometric determination of gMFI), was then assessed by supernatant analysis or intracellular staining. n = 3–16 replicates per group; N = 3 independent experiments.

**Table S1**

**Table S1.** Oligonucleotide sequences used in this study.

| Oligonucleotide | Sequence (5′→3′) |
|---|---|
| ssRNA | CAG CUC UAG A-Cy3 |
| cRNA | UCU AGA GCU G |
| ssDNA | CAG CTC TAG A-Cy3 |
| cDNA | TCT AGA GCT G |
| (rA)10 | (rA)10-Cy3 |
| (rC)10 | (rC)10-Cy3 |
| (rU)10 | (rU)10-Cy3 |
| (rGA)5 | (rGA)5-Cy3 |
| 10 nt-ATTO488N with two U | CAG CUC UAG A /3ATTO488N/ |
| 10 nt-ATTO647N with two U | CAG CUC UAG A /3ATTO647N/ |
| (dU)10-Cy3Sp | (dU)10-Cy3Sp |
| (dA)10-Cy3Sp | (dA)10-Cy3Sp |
| (dC)10-Cy3Sp | (dC)10-Cy3Sp |
| (dGdA)5-Cy3Sp | (dGdA)5-Cy3Sp |

### Table S2

**Table S2.** Statistical analysis of exosome uptake (fluorescence intensity per cell) across five experimental conditions for Figure 4e. Individual uptake values were analyzed without excluding any non-zero data points. Because the raw distributions were highly right-skewed and contained true zeros, data were transformed using log1p(x) = ln(1 + YEx) prior to statistical testing. A one-way ANOVA was performed on the log-transformed values ($F(4, 473) = 14.74$, $p = 2.34\times10^{-11}$), followed by Tukey's Honest Significant Difference (HSD) test for all pairwise comparisons with multiplicity-adjusted p-values reported. Reported differences therefore represent adjusted mean separations on the log scale.

| Group1 | Group2 | Mean difference of log1p | Adjusted p value | 95% CI lower | 95% CI upper | Significant |
|---|---|---|---|---|---|---|
| No Exo | Exo | −1.043 | $1.53 \times 10^{-5}$ | −1.567 | −0.520 | Yes |
| No Exo | Ruc-Exo | −1.757 | $1.51 \times 10^{-7}$ | −2.231 | −1.283 | Yes |
| Exo | Ruc-Exo | −0.714 | $6.84 \times 10^{-4}$ | −1.148 | −0.279 | Yes |
| Ruc-Exo | Ruc-Exo + DNase I | 1.748 | $2.25 \times 10^{-5}$ | 1.225 | 2.271 | Yes |
| Ruc-Exo | Ruc-Exo + mBSA | 0.723 | $2.91 \times 10^{-4}$ | 0.260 | 1.186 | Yes |
| No Exo | Ruc-Exo + DNase I | −0.009 | $9.89 \times 10^{-1}$ | −0.533 | 0.515 | No |
| No Exo | Ruc-Exo + mBSA | −1.034 | $2.77 \times 10^{-5}$ | −1.548 | −0.521 | Yes |
| Exo | Ruc-Exo + DNase I | 1.034 | $9.79 \times 10^{-1}$ | 0.501 | 1.568 | No |
| Exo | Ruc-Exo + mBSA | 0.009 | $9.65 \times 10^{-1}$ | −0.532 | 0.550 | No |
| Ruc-Exo + DNase I | Ruc-Exo + mBSA | −1.025 | $9.32 \times 10^{-1}$ | −1.575 | 0.475 | No |

**Table S3**

**Table S3.** Statistical analysis of exosome uptake (particles per cell) across five experimental conditions for Figure S35. Individual uptake values were analyzed without excluding any non-zero data points. Because the raw distributions were highly right-skewed and contained true zeros, data were transformed using log1p(x) = ln(1 + x) prior to statistical testing. A one-way ANOVA was performed on the log-transformed values ($F(4, 429) = 93.71$, $p = 3.21 \times 10^{-57}$), followed by Tukey's Honest Significant Difference (HSD) test for all pairwise comparisons with multiplicity-adjusted p-values reported. Reported differences therefore represent adjusted mean separations on the log scale.

| Group1 | Group2 | Mean difference of log1p | Adjusted p value | 95% CI lower | 95% CI upper | Significant |
|---|---|---|---|---|---|---|
| **No Exo** | Exo | −0.407 | $1.34 \times 10^{-8}$ | −0.523 | −0.291 | Yes |
| **No Exo** | Ruc-Exo | −1.296 | $2.52 \times 10^{-57}$ | −1.394 | −1.198 | Yes |
| **Exo** | Ruc-Exo | −0.889 | $1.46 \times 10^{-43}$ | −0.994 | −0.784 | Yes |
| **Ruc-Exo** | Ruc-Exo + DNase I | 0.776 | $3.12 \times 10^{-43}$ | 0.664 | 0.888 | Yes |
| **Ruc-Exo** | Ruc-Exo + mBSA | 0.678 | $2.66 \times 10^{-34}$ | 0.560 | 0.796 | Yes |
| **No Exo** | Ruc-Exo + DNase I | −0.520 | $3.24 \times 10^{-14}$ | −0.654 | −0.386 | Yes |
| **No Exo** | Ruc-Exo + mBSA | −0.618 | $2.22 \times 10^{-20}$ | −0.748 | −0.488 | Yes |
| **Exo** | Ruc-Exo + DNase I | −0.113 | $2.40 \times 10^{-1}$ | −0.279 | 0.053 | No |
| **Exo** | Ruc-Exo + mBSA | −0.211 | $1.58 \times 10^{-3}$ | −0.374 | −0.048 | Yes |
| **Ruc-Exo + DNase I** | Ruc-Exo + mBSA | −0.098 | $4.19 \times 10^{-1}$ | −0.249 | 0.052 | No |
